# Glial Ca^2+^ Signaling Links Endocytosis to K^+^ Buffering around Neuronal Somas to Regulate Excitability

**DOI:** 10.1101/426130

**Authors:** Shirley Weiss, Jan E. Melom, Kiel G. Ormerod, Yao V. Zhang, J. Troy Littleton

## Abstract

Glial-neuronal signaling at synapses is widely studied, but how glia interact with neuronal somas to regulate neuronal function is unclear. *Drosophila* cortex glia are restricted to brain regions devoid of synapses, providing an opportunity to characterize interactions with neuronal somas. Mutations in the cortex glial *NCKX*^*zydeco*^ elevate basal Ca^2+^, predisposing animals to seizures. To determine how cortex glial Ca^2+^ signaling controls neuronal excitability, we performed an *in vivo* modifier screen of the *NCKX*^*zydeco*^ seizure phenotype. We show that elevation of glial Ca^2+^ causes hyperactivation of calcineurin-dependent endocytosis and accumulation of early endosomes. Knockdown of sandman, a K_2P_ channel, recapitulates *NCKX*^*zydeco*^ seizures. In addition, restoring glial K^+^ buffering by ectopically expressing a leak K^+^ channel abolishes *NCKX*^*zydeco*^ seizures. These data provide an unexpected link between glial Ca^2+^ signaling and the more well-known roles of glia in K^+^ buffering as a key mechanism for regulating neuronal excitability.

**Impact statement:** The current study provides evidence for a direct link between glial calcium signaling to classical functions of glia in buffering external K^+^ as a mechanism to regulate neuronal excitability.

## Introduction

Glial cells are well known to play structural and supportive roles for their more electrically excitable neuronal counterparts. However, growing evidence indicates glial Ca^2+^ signaling influences neuronal physiology on a rapid time scale. In the cortex, glia and neurons exist in equal abundance (Azevedo et al., 2009) and are intimately associated. A single astrocytic glia contacts multiple neuronal cell bodies, hundreds of neuronal processes, and tens of thousands of synapses (Halassa et al., 2007; Ventura and Harris, 1999). Cultured astrocytes oscillate intracellular Ca^2+^ spontaneously (Takata and Hirase, 2008) and in response to neurotransmitters (Agulhon et al., 2008; Lee et al., 2010), including glutamate (Cornell-Bell et al., 1990). Glutamate released during normal synaptic transmission is sufficient to induce astrocytic Ca^2+^ oscillations (Dani et al., 1992; Wang et al., 2006), which trigger Ca^2+^ elevation in co-cultured neurons (Nedergaard, 1994; Parpura et al., 1994) that can elicit action potentials (Angulo et al., 2004; Fellin et al., 2006; Fellin et al., 2004; Pirttimaki et al., 2011). These astrocyte-neuron interactions suggest abnormally elevated glial Ca^2+^ might produce neuronal hypersynchrony. Indeed, increased glial activity is associated with abnormal neuronal excitability (Wetherington et al., 2008), and pathologic elevation of glial Ca^2+^ can play an important role in the generation of seizures (Gomez-Gonzalo et al., 2010; Tian et al., 2005). However, the molecular pathway(s) by which glia-to-neuron communication alters neuronal excitability is poorly characterized. In addition, how glia interface with synaptic versus non-synaptic regions of neurons is unclear.

Several glia-neuronal cell body interactions have been reported for different glial subtypes (Allen and Barres, 2009; Baalman et al., 2015; Battefeld et al., 2016; Takasaki et al., 2010). A single mammalian astrocyte can be associated with multiple neuronal cell bodies and thousands of synapses (Halassa et al., 2007; Ventura and Harris, 1999). However, the complex structure of mammalian astrocytes and the diversity of their glia-neuron contacts makes it challenging to directly manipulate glial signaling only at contacts with neuronal cell bodies. *Drosophila* provides an ideal system to study glial-neuronal soma interactions as the *Drosophila* CNS contains two specialized astrocyte-like glial subtypes that interact specifically either with dendrites and synapses (astrocytes, Stork et al., 2014) or with neuronal cell bodies (cortex glia, Awasaki et al., 2008; Pereanu et al., 2005). Cortex glia processes encapsulate all neuronal cell bodies in the CNS with fine, lattice like processes (Awasaki et al., 2008; Coutinho-Budd et al., 2017) (Fig. S1A), and are thought to provide metabolic support and electrical isolation to their neuronal counterparts (Buchanan and Benzer, 1993; Volkenhoff et al., 2015).

Previous work in our lab identified zydeco (zyd), a cortex glial enriched Na^+^Ca^2+^K^+^ (NCKX) exchanger involved in maintaining normal neural excitability (Melom and Littleton, 2013). Mutations in *NCKX*^*zydeco*^ (hereafter referred to as *zyd*) predispose animals to temperature-sensitive seizures and result in bang sensitivity (seizures induced following a brief vortex). Basal intracellular Ca^2+^ levels are elevated in *zyd* cortex glia, while near-membrane microdomain Ca^2+^ oscillations observed in wildtype cortex glia are abolished. Whether the loss of Ca^2+^ microdomain events in *zyd* is due to a disruption in the mechanism generating these events or secondary to a saturation effect from elevated basal Ca^2+^ levels is unclear. Though the mechanism(s) by which cortex glia modulate neuronal activity in *zyd* mutants remains unclear, disruption of glial Ca^2+^ regulation dramatically enhances seizure susceptibility.

To determine how altered cortex glial Ca^2+^ signaling in *zyd* mutants regulates neuronal excitability, we took advantage of the *zyd* mutation and performed an RNAi screen for modifiers of the seizure phenotype. Here we show that chronic elevation of glial Ca^2+^ causes hyperactivation of calcineurin-dependent endocytosis leading to an endo-exocytosis imbalance. In addition, knockdown of sandman, a K_2P_ channel, recapitulates *zyd* seizures and acts downstream of calcineurin in cortex glia. Sandman was previously identified as the key K^+^ channel that cycles to and from the plasma membrane in a group of central complex neurons to modulate their excitability and control sleep homeostasis in the *Drosophila* circadian pathway (Pimentel et al., 2016). Our findings suggest similar regulation of sandman in cortex glia could allow dynamic control of K^+^ levels surrounding neuronal somas as a mechanism to gate neuronal excitability. Indeed, overexpression of a constitutively active K^+^ channel in cortex glia can rescue *zyd* seizures. Together, these findings suggest glial Ca^2+^ interfaces with calcineurin-dependent endocytosis to regulate plasma membrane protein levels and the K^+^ buffering capacity of glia associated with neuronal somas. Disruption of these pathways leads to enhanced neuronal excitability and seizures, suggesting potential targets for future glial-based therapeutic modifiers of epilepsy.

## Results

### Mutations in a cortex glial NCKX generate stress-induced seizures without affecting brain structure or baseline neuronal function

We previously identified and characterized a *Drosophila* temperature-sensitive (TS) mutant termed *zydeco* (*zyd*) that exhibits seizures when exposed to a variety of environmental stressors, including heat-shock and acute vortex (Melom and Littleton, 2013). The *zyd* mutation maps to a NCKX exchanger that extrudes cytosolic Ca^2+^. Restoring zyd function specifically in cortex glial completely reverses the *zyd* seizure phenotype. Cortex glia exhibit spatial segregation reminiscent of mammalian astrocytes, with each glial cell ensheathing multiple neuronal somas (Awasaki et al., 2008; Melom and Littleton, 2013). However, little is known about their role in the mature nervous system. *In vivo* Ca^2+^ imaging using a myristoylated Ca^2+^ sensitive-GFP (myrGCaMP5) revealed small, rapid cortex glial Ca^2+^ oscillations in wildtype *Drosophila* larvae. In contrast, *zyd* mutants lack microdomain Ca^2+^ transients and exhibit elevated baseline intracellular Ca^2+^, indicating altered glial Ca^2+^ regulation underlies seizure susceptibility in *zyd* mutants (Melom and Littleton, 2013).

Given cortex glia regulate the guidance of secondary axons and maintenance of cortical structural integrity (Coutinho-Budd et al., 2017; Dumstrei et al., 2003; Spindler et al., 2009), we first tested whether the *zyd* TS seizure phenotype might arise secondary to developmental changes or defective assembly of brain circuits. We knocked down ZYD chronically throughout development or conditionally in adult stages following brain development using a UAS-zyd^RNAi^ hairpin expressed with a pan-glial driver (repo-gal4) or cortex glial-specific drivers (NP2222-gal4 and GMR54H02-gal4). Though *zyd* mutations were shown to be viable (Guan et al., 2005; Melom and Littleton, 2013), chronic pan-glial knock down of zyd caused lethality, with >95% of animals dying before the 3^rd^ instar larval stage (Fig. S1B). This difference might arise from RNAi knockdown of zyd leading to a functional null, or from the requirement of zyd in glial subtypes other than cortex glia, while it is only required in cortex glia for the generation of seizures (given that cortex glial specific over expression of zyd can completely rescue *zyd* seizure phenotype (Melom and Littleton, 2013)). In contrast, both chronic and inducible cortex glial knockdowns mimicked the *zyd* TS seizure phenotype (Figs. 1A), suggesting the *zyd* TS phenotype does not arise from a developmental defect. Seizure characteristics, including temperature threshold for seizure initiation and seizure kinetics, were similar between 3^rd^ instar larvae and adults (Fig. 1B-C), indicating a comparable requirement for ZYD at both stages. Morphological examination of brain structure and cortex glial morphology using fluorescent microscopy revealed no apparent changes in *zyd* mutants (Fig. S1C). To determine if loss of ZYD affected glial or neuronal cell survival, we quantified the cell death marker DCP-1 (cleaved death caspase protein-1 (Akagawa et al., 2015)) in control and *zyd* 3^rd^ instar larvae and adults. No change in cleaved DCP1 levels were found, indicating basal cell death was unaffected (Fig. S1D). Together, these results suggest that the *zyd* TS seizure phenotype is not due to morphological or developmental changes in brain anatomy, or changes in the ability of cortex glia to ensheath neuronal cell bodies.

**Figure 1.**
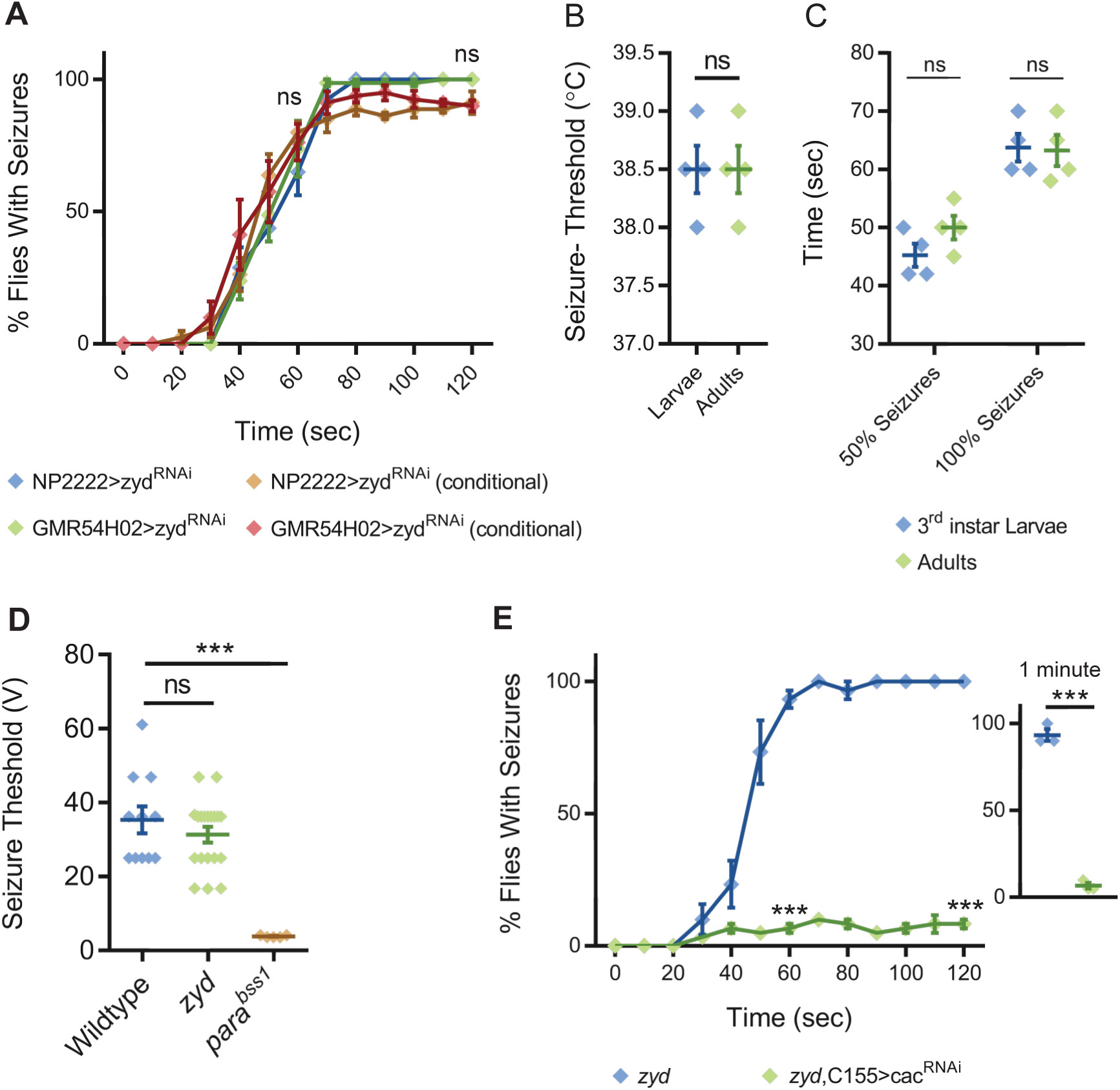
Mutations in a cortex glial NCKX generate stress-induced seizures. **A.** Time course of HS-induced seizures (38.5°C, HS) following chronic or conditional knockdown of zyd with two different cortex glial drivers (NP2222 and GMR54H02) is shown. Rearing adult flies at the restrictive temperature (>30°C) for gal80^ts^ (a temperature-sensitive form of the gal4 inhibitor, gal80, see methods) removes gal80 inhibition of gal4 and allows expression of zyd^RNAi^ only at the adult stage. These manipulations reproduce the *zyd* mutant seizure phenotype (N=4 groups of 20 flies/genotype). **B-C.** Behavioral analysis of HS-induced seizures at 38.5°C shows that larval and adult seizures have similar temperature threshold (**B**) and kinetics (**C**) (N=4 groups of 10-20 animals/condition/treatment). **D.** Giant fiber recordings of seizure threshold in wildtype, *zyd* and Para^bss1^ (positive control) are shown. The voltage required to induce seizures in *zyd* is not significantly different from wildtype (35.32 ± 3.65V and 31.33 ± 2.12V, p=0.3191, n≥7 flies/genotype). **E.** Behavioral analysis of the time course of HS-induced seizures indicates neuronal knockdown of cac (C155>cac^RNAi^) rescues the *zyd* seizure phenotype. Inset shows results after 1 minute of HS (p=0.0004, N=4 groups of 20 flies/genotype). Error bars are SEM, ***=P<0.001, Student’s t-test.

NCKX transporters use the gradients of Na^+^ and K^+^ ions to extrude Ca^2+^ out of cells, and it is possible that the loss of ZYD would not only cause an elevation of intracellular Ca^2+^ (Melom and Littleton, 2013), but will also change the ionic balance of other ions (i.e. Na^+^ and K^+^). This in turn might directly affect the intrinsic membrane properties of surrounding neurons and their seizure susceptibility, and possibly other circuits mediating specific behaviors. Hence, we assayed if *zyd* animals display altered behaviors or neuronal excitability in the absence of the trigger needed to induce seizures. We used a gentle touch assay (Ma et al., 2016; Zhou et al., 2012) to investigate whether the *zyd* mutation changes larval startle-induced behaviors, as elevated Ca^2+^ activity in astrocytes was reported to correlate with elevated arousal in mice (Ding et al., 2013; Paukert et al., 2014; Srinivasan et al., 2015) and in *Drosophila* (Ma et al., 2016). Crawling 3^rd^ instar larvae touched anteriorly execute one of two responses: pausing and/ or continuing forward (type I response) or an escape response by crawling backwards (type II response), considered to be the larval startle-induced behavior. We found that wildtype, *zyd* and NP2222>zyd^RNAi^ larvae exhibited similar frequencies of type I and type II responses (Fig. S1E). In addition, adult *zyd* flies exhibited normal locomotion (Fig. S1F) and larvae exhibited normal light avoidance responses at room temperature (Fig. S1G), indicating baseline neuronal properties required for these behaviors are unaffected.

To directly measure changes in neuronal excitability that might contribute to *zyd* seizures, we performed electrophysiological recordings in the giant fiber system (GFS, (Pavlidis and Tanouye, 1995)). In this assay, GF neurons are stimulated in the adult brain and the output response is recorded. By progressively increasing stimulation intensity, the voltage threshold that triggers seizure activity in the GF circuit can be directly assayed, providing a readout of neuronal excitability. Most *Drosophila* bang-sensitive mutations display seizure induction at much lower voltages, consistent with their primary effect on membrane excitability thresholds within the neuron. For example, *para*^*bss1*^, a gain-of-function bang sensitive mutation in the voltage-gated Na^+^ channel, dramatically lowers seizure threshold (Fig. 1D) (Parker et al., 2011). In contrast, the voltage threshold for seizure induction in *zyd* was not significantly different from controls (Fig. 1D), indicating basic intrinsic neuronal properties are not altered in *zyd* animals at rest. Our previous finding that cortex glial knockdown of calmodulin (cam, (Melom and Littleton, 2013)) can completely reverse *zyd* seizure phenotype, further support the notion that glial regulation of neuronal excitability in *zyd* flies proceeds through a glial calmodulin-dependent pathway, and is not simply due to changes in ionic balance leading to altered intrinsic neuronal properties.

To determine if elevated neuronal activity is required for the stress-induced seizures in *zyd* mutants, we assayed seizure behavior in animals with reduced synaptic transmission and neuronal activity. Pan-neuronal knockdown of Cacophony (cac), the presynaptic voltage-gated Ca^2+^ channel responsible for neurotransmitter release (Kawasaki et al., 2004; Rieckhof et al., 2003), significantly reduced neuronal activity (Fig. S1H) and rescued *zyd* TS-induced seizures (Fig. 1E). These findings indicate elevated neuronal activity in *zyd* mutants during the heat shock is required for seizure induction following dysregulation of cortex glial Ca^2+^.

### A genetic modifier screen of the zyd seizure phenotype reveals glia to neuron signaling mechanisms

To elucidate pathways by which cortex glial Ca^2+^ signaling controls somatic regulation of neuronal function and seizure susceptibility, we performed a targeted RNAi screen for modifiers of the *zyd* TS seizure phenotype in adult animals. We reasoned that removal of a gene product required for this signaling pathway would prevent *zyd* TS seizures when absent. We used the pan-glial driver repo-gal4 to express RNAi to knockdown 847 genes encoding membrane receptors, secreted ligands, ion channels and transporters, vesicular trafficking proteins and known cellular Ca^2+^ homeostasis and Ca^2+^ signaling pathway components (Tables S1, S2). Given the broad role of Ca^2+^ as a regulator of intracellular biology, we expected elevated Ca^2+^ levels in *zyd* mutants to interface with several potential glial-neuronal signaling mechanisms. Indeed, the screen revealed multiple genetic interactions, identifying gene knockdowns that completely (28) or partially (21) rescued *zyd* seizures, caused lethality on their own (95) or synthetic lethality in the presence of the *zyd* mutation (5), enhanced *zyd* seizures (37) or triggered seizures (3) in a wildtype background (Table S1).

Given TS seizures in *zyd* mutants can be fully rescued by reintroducing wildtype ZYD in cortex glia, we expected the genes identified in the RNAi pan-glial knockdown screen to function specifically within this population of cells. To directly examine cell-type specificity of the suppressor hits, we knocked down the top 33 rescue RNAis with cortex glial specific drivers (NP2222-gal4 and GMR54H02-gal4, Table S1). For the majority of suppressors, rescue with cortex glial specific drivers was weaker, either due to lower RNAi expression levels compared to the stronger repo-gal4 driver or due to the requirement of the gene in other glial subtypes as well. To validate the rescue effects we observed, non-overlapping RNAis or mutant alleles for these genes were also assayed. For the current analysis, we focused only on the characterization of cortex glial Ca^2+^-dependent pathways that are mis-regulated in *zyd*, and how this mis-regulation promotes neuronal seizure susceptibility.

### Cortex glial calcineurin activity is required for seizures in zyd mutants

We previously observed that knockdown of glial calmodulin (cam) eliminates the *zyd* seizure phenotype (Melom and Littleton, 2013), suggesting a Ca^2+^/cam-dependent signaling pathway regulates glial to neuronal communication in *zyd*. Cam is an essential Ca^2+^-binding protein that regulates multiple Ca^2+^-dependent cellular processes and is abundantly expressed in *Drosophila* glia (Altenhein et al., 2006), although its role in glial biology is unknown. In the RNAi screen for *zyd* interactors, pan-glial knockdown of the regulatory calcineurin (CN) B subunit, CanB2, completely rescued both heat-shock and vortex induced seizures in *zyd* animals (Fig. 2A-B). Recordings of motor central pattern generator (CPG) output at the larval neuromuscular junction (NMJ), showed that in contrast to the continuous neuronal firing observed in *zyd* mutants, recordings from *zyd*;;repo>CnB2^RNAi#1^ larvae exhibit normal rhythmic firing at 38°C similar to wildtype controls (Fig. 2B). Calcineurin (CN) is a highly conserved Ca^2+^/cam-dependent protein phosphatase implicated in a number of cellular processes in mammals (Rusnak and Mertz, 2000). The rescue effect of CanB2 knockdown on *zyd* seizure phenotype was similar when CanB2 was targeted using three additional, non-overlapping CanB2 RNAi constructs (Fig. 2A, 2F). Knocking down CanB2 on a wildtype background was viable (Fig. S2A) and did not cause any significant change in larval light avoidance (Fig. S2B) or adult locomotion and activity (Fig. S2C). To refine the glial subpopulation in which CanB2 activity is necessary to promote seizures in *zyd* mutants, we knocked down CanB2 using cortex glial specific drivers. CanB2 knockdown in astrocytes resulted in no rescue of the *zyd* phenotype (Fig. 2D, S2D). CanB2 knockdown with a cortex-glial specific driver (NP2222-gal4) greatly improved the *zyd* phenotype. Animals no longer displayed continuous seizures, but the rescue was less robust compared to pan-glial knockdown (Fig. 2D, Fig. S2D), possibly due to lower expression level of the RNAi. Indeed, rescue was greatly enhanced by cortex glial-specific knockdown of CanB2 using two copies of the CanB2 RNAi construct (Fig. 2D), with ∼90% of *zyd*;NP2222>CanB2^2xRNAi^ adult animals lacking seizures. The effect of CanB2 knockdown was specific to cortex glia, as it was insensitive to blockade of expression of CanB2^RNAi^ in neurons using C155–gal80 (a neuron specific gal4 repressor; Fig. 2D and S2D). To exclude a developmental effect of CanB2^RNAi^ knockdown within glia, we blocked gal4/UAS-driven CanB2^RNAi^ with gal80^ts^ and expressed a single copy of CanB2^RNAi^ only in adult *zyd* mutants. Adult flies reared at the permissive temperature for gal80^ts^ to allow CanB2^RNAi^ expression exhibited significantly less seizures after 3 days, with only ∼20% of flies displaying *zyd*-like seizures by 5 days (Fig. 2E). *Zyd* seizure rescue by CanB2 knockdown did not result from simple alterations in motility, as repo>CanB2^RNAi^ and *zyd*;;repo>CanB2^RNAi^ animal exhibited normal larval light avoidance responses (Fig. S2E) and adult locomotion (Fig. S2F). We conclude that CanB2 is required in cortex glia to promote *zyd* TS seizure activity.

**Figure 2.**
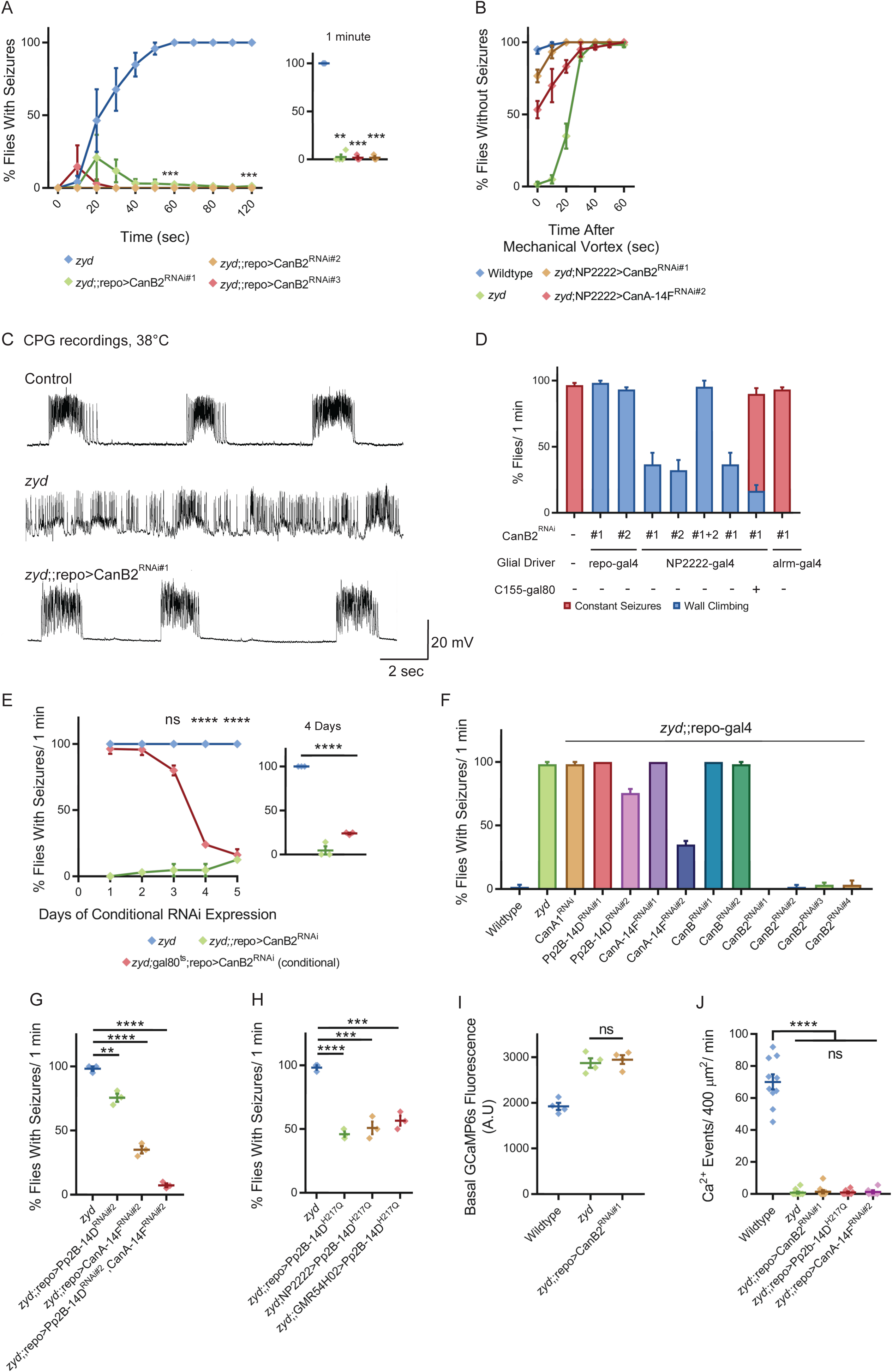
Cortex glial knockdown of calcineurin rescues *zyd* seizures without affecting intracellular Ca^2+^. **A.** Behavioral analysis of HS-induced seizures. Pan-glial knockdown of the CN regulatory subunit, CanB2, with three different hairpins (#1, #2 and #3, see methods) rescues the *zyd* seizure phenotype (N=4 groups of ≥15 flies/genotype). Inset shows analysis after 1 minute of HS (p=0.0001). **B.** Behavioral analysis of the recovery from vortex-induced seizures. Pan-glial knockdown of CanB2 and CanA-14F rescues *zyd* vortex-induced seizures (N=3 groups of 20 flies/genotype). **C.** Representative voltage traces of spontaneous CPG activity at larval 3^rd^ instar muscle 6 at 38°C in wildtype, *zyd* and *zyd*;;repo>CanB2^RNAi#1^ animals (*n*≥5 preparations/genotype). **D.** Detailed analysis (see methods) of HS induced behaviors of *zyd*/CanB2^RNAi^ flies. Cortex glial knockdown of CanB2 leads to seizure rescue in ∼30% of *zyd*;NP2222>CanB2^RNAi^ flies, with the remaining ∼70% displaying partial rescue. Cortex glial CanB2 knockdown with two copies of the RNAi (*zyd*;NP2222>CanB2^2xRNAi^) recapitulates the full rescue seen with pan-glial knockdown. Inhibiting gal4 expression of the RNAi in neurons with gal80 (C155-gal80) does not alter the rescue observed with cortex glial knockdown, and astrocyte specific (alrm-gal4) CanB2 knockdown does not rescue *zyd* seizures (N=3 groups of >15 flies/genotype, see supplemental Fig. S2B for complete dataset). **E.** Cortex glial conditional knockdown of CanB2 using gal4/gal80^ts^. Rearing adult flies at the restrictive temperature (>30°C) for gal80^ts^ allows expression of CanB2^RNAi^ only at the adult stage. A significant reduction in seizures (p<0.0001) was seen after four days of rearing flies at the restrictive temperature for gal80^ts^ (31°C), with only ∼25% of adults showing seizures. The reduction in seizures was enhanced when adults were incubated at 31°C for longer periods (N=3 groups of >10 flies/genotype). Inset shows analysis after 4 days of incubation at 31°C (p=0.0001). **F.** Pan-glial knockdown of the *Drosophila* calcineurin (CN) family (CanA1, CanA-14D/ Pp2B-14D, CanA-14F, CanB and CanB2) indicate CanB2 knockdown completely rescues *zyd* seizures, CanA14D and CanA14F knockdowns partially reduce seizures, and knockdown of either CanA1 or CanB do not alter the *zyd* phenotype (N=4 groups of >10 flies/genotype). **G.** Pan-glial knockdown of CanA14D and CanA14F partially rescues the *zyd* HS seizures phenotype (∼25% rescue for CanA14D, p=0.0032; and ∼60% rescue for CanA14F, p<0.0001). Knocking down the two genes simultaneously rescues *zyd* seizures, with only ∼10% of flies showing seizures (∼90% rescue, p<0.0001, N=3 groups of >10 flies/genotype). **H.** Overexpressing a dominant-negative form on Pp2B-14D (CanA^H217Q^) rescues ∼50% of *zyd* seizures regardless of the driver used (repo: p<0.0001; NP2222: p=0.0006; GMR54H02-p=0.0004. N=3 groups of >10 flies/ genotype). **I.** Larval Ca^2+^ imaging in cortex glia expressing myrGCaMP6s indicates the elevated basal Ca^2+^ fluorescence at 25°C observed in *zyd* mutants relative to wildtype cortex glia (p=0.0003) is not altered following CanB2 knockdown (*zyd*;;repo>CanB2^RNAi^, p=0.6096. n≥5 animals/genotype). **J.** Microdomain Ca^2+^ oscillations observed in wildtype cortex glia expressing myrGCaMP6s are abolished in *zyd* cortex glia and are not restored following either CanB2 or CanA14F knockdown (n≥5 animals/genotype). Error bars are SEM, **=P<0.01, ***=P<0.001, ****=P<0.0001, Student’s t-test.

CN is a heterodimer composed of a ∼60 kDa catalytic subunit (CanA) and a ∼19 kDa EF-hand Ca^2+^-binding regulatory subunit (CanB). Both subunits are essential for CN phosphatase activity. The *Drosophila* CN gene family contains three genes encoding CanA (CanA1, Pp2B-14D and CanA-14F) and two genes encoding CanB (CanB and CanB2) (Takeo et al., 2006). Previous studies in *Drosophila* demonstrated several CN subunits (CanA-14F, CanB, and CanB2) are broadly expressed in the adult *Drosophila* brain (Tomita et al., 2011) and that neuronal CN is essential for regulating sleep (Nakai et al., 2011; Tomita et al., 2011). CN function within glia has not been characterized. We found that pan-glial knockdown of two CanA subunits, Pp2B-14D and CanA*-*14F, partially rescued *zyd* heat-shock and vortex-induced seizures (Fig. 2B, F). The rescue was more robust for vortex-induced seizures than those induced by heat-shock, suggesting heat-shock is likely to be a more severe hyperexcitability trigger. Rescue was enhanced by knockdown of both Pp2B-14D and CanA*-*14F (Fig. 2G), with more than ∼90% of *zyd*;;repo>2xCanA^RNAi^ flies lacking seizures, suggesting a redundant function of these two subunits in glial cells. Similar to CanB2, knockdown of both Pp2b-14D and CanA-14F on a wildtype background was viable (Fig. S2A) and did not cause any significant change in larval light avoidance (Fig. S2B) or adult locomotion and activity (Fig. S2C). To further verify the effect of CanA knockdown and the glial subpopulation in which CN activity is required, we overexpressed a dominant negative (DN) form of CanA (Pp2B-14D^H217Q^) using either pan-glial (repo-gal4) or two different cortex-glial specific drivers (NP2222-gal4 and GMR54H02-gal4). Overexpressing Pp2B-14D^H217Q^ resulted in ∼50% of *zyd*;;Pp2B-14D^H217Q^ flies becoming seizure-resistant, regardless of the driver used (Fig 2H). Imaging of intracellular Ca^2+^ in cortex glia with GMR54H02-gal4 driving myrGCaMP6s revealed that CN knockdown had no effect on either the elevated basal Ca^2+^ levels or the lack of microdomain Ca^2+^ events observed in *zyd* cortex glia (Fig. 2I-J). These observations indicate CN function is required downstream of elevated intracellular Ca^2+^, rather than to regulate Ca^2+^ influx or efflux in cortex glial cells. Together, these results demonstrate that a CN-dependent signaling mechanism in cortex glia is required for a glia to neuronal pathway that drives seizure generation in *zyd* mutants.

### Calcineurin activity is enhanced in zyd cortex glia

To characterize CN activity in wildtype and *zyd* cortex glia we used the CalexA system as a reporter for CN activity. CalexA (Ca^2+^-dependent nuclear import of LexA) (Masuyama et al., 2012) was originally designed for labeling active neurons in behaving animals. In this system, sustained neural activity induces CN activation and dephosphorylation of a chimeric transcription factor LexA-VP16-NFAT (termed CalexA) which is then transported into the nucleus. The imported dephosphorylated CalexA drives GFP reporter expression in active neurons (for schematic representation, see Fig. S2G). The CalexA components were brought into control and *zyd* mutant backgrounds to directly assay CN activity. A substantial basal activation of CN was observed in control 3^rd^ instar larval cortex glia at room temperature using fluorescent imaging (Fig. 3A). CN activity and the resulting GFP expression was enhanced in *zyd* cortex glia (Fig. 3B) and greatly reduced in *zyd*/CanB2^RNAi^ cortex glia (Fig. 3C). Western blot analysis of CalexA-induced GFP expression in adult head extracts revealed enhanced cortex glial CN activity in adult *zyd* mutants compared to controls as well (23 ± 3% enhancement, Fig 3D-E). RNAi knockdown of CanB2 greatly reduced CalexA GFP expression as expected (27 ± 1% reduction, Fig. 3C-E). These results demonstrate CN activity is enhanced downstream of the elevated Ca^2+^ levels in *zyd* mutant cortex glia, and that CN activity can be efficiently reduced by RNAi knockdown of CanB2.

**Figure 3.**
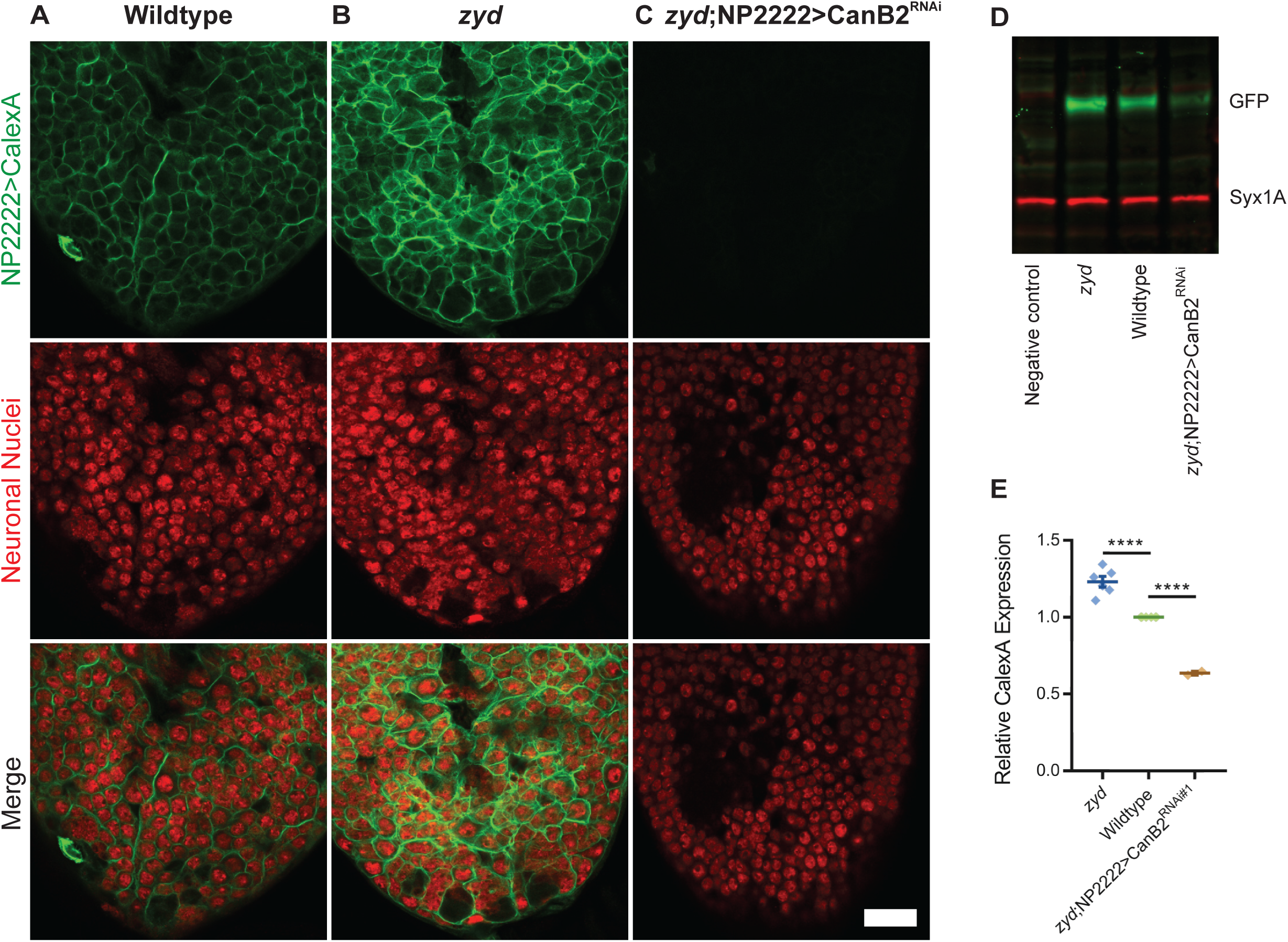
Calcineurin activity is enhanced in *zyd* cortex glia and can be efficiently suppressed by CanB2 knockdown. **A-C.** Fluorescence microscopy imaging of cortex glial CalexA-derived GFP expression in wildtype (**A**), *zyd* (**B**) and *zyd*;NP2222>CanB2^RNAi^ (**C**) larvae. Red: anti-elav=neuronal nuclei; green: anti-GFP=cortex glial CN activity (animals were reared at 25°C, Scalebar = 20 μm, N≥5 animals/ genotype). **D-E.** Western blot analysis of cortex glial CalexA derived (NP2222>CalexA) GFP expression in *zyd*, wildtype and *zyd*;NP2222>CanB2^RNAi^ adult heads. CN activity is enhanced by ∼25% (p<0.0001) in *zyd* cortex glia and reduced by ∼35% (p<0.0001) in CanB2^RNAi^ animals (N≥2 experiment, 5 heads/ sample). GFP signals in each experiment were normalized to wildtype. Error bars are SEM, ****=P<0.0001, Student’s t-test.

### Pharmacological targeting of the glial calcineurin pathway rescues zyd seizures

Several seizure mutants in *Drosophila* can be suppressed by commonly used anti-epileptic drugs (Kuebler and Tanouye, 2002; Song and Tanouye, 2008), indicating conservation of key mechanisms that regulate neuronal excitability. CN activity is strictly controlled by Ca^2+^ levels, calmodulin, and CanB, and can be inhibited by the immunosuppressants cyclosporine A (CsA) and FK506. The CN inhibitor FK506 has been previously shown to reduce seizures in a rodent kindling model (Moia et al., 1994; Moriwaki et al., 1996), suggesting CN can modulate epilepsy in mammals. To assay if *zyd* TS seizures can be prevented with an anti-CN drug, adult flies were fed with media containing CsA and tested for HS induced seizures after 0, 3, 6, 12 and 24 hours of drug feeding (red arrowheads in Fig. 4A, Fig. 4B-C). *Zyd* flies fed with 1 mM CsA for 12 hours showed ∼80% less seizures than controls (Fig. 4B, C). Feeding the flies for longer periods or with higher dosages of CsA led to changes in basal activity and lethality, and hence flies were not tested for their HS seizure behavior beyond this point. Indeed, high-dose feeding of flies was previously reported to result in lethality (Lee et al., 2016). Seizure rescue by CsA was dose-dependent, with less robust suppression when flies were fed with 0.3 mM CsA (Fig. 4B, D). The CsA rescue was reversible, as *zyd* seizures reoccurred following 12 hours of CsA withdrawal (Fig. 4C). Although the mechanism of action of CsA primarily involves inhibiting the function of CN, it is also predicted to have additional targets including calpain2 and p38 (Hu et al., 2014). However, pan-glial or cortex glial-specific knockdown of the *Drosophila* calpain (calpain 1, calpain 2 and calpain 3) and p38 (p38a, p38b an p38c) gene families did not result in any behavioral change in *zyd* nor wildtype flies. We conclude that pharmacologically targeting the glial CN pathway can improve the outcome of glial-derived neuronal seizures in the *zyd* mutant.

**Figure 4.**
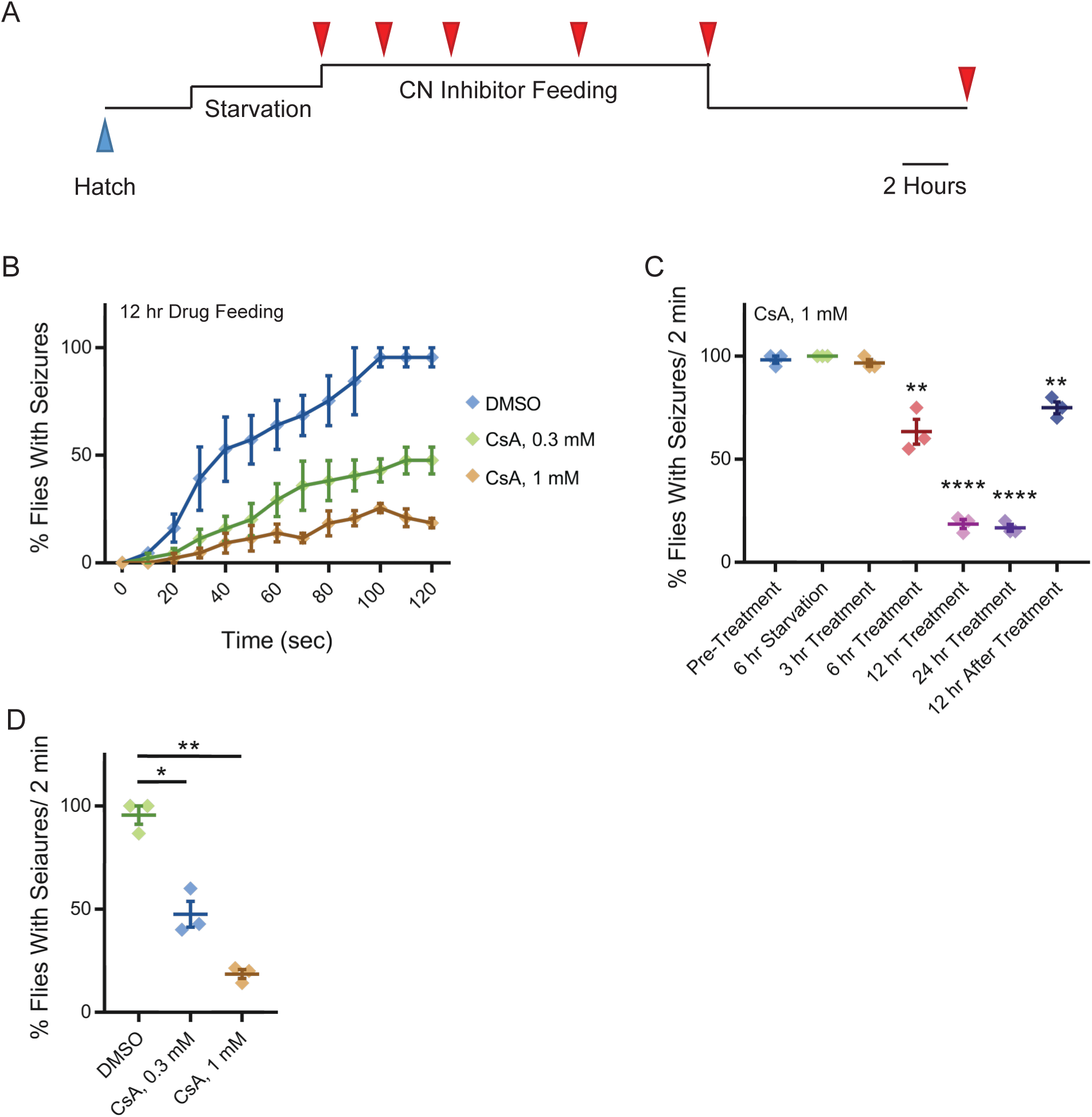
Pharmacologically targeting calcineurin activity suppresses *zyd* heat shock-induced seizures. **A.** Schematic representation of the experimental design. Adult male flies (<1 day old) were starved for 6 hours and fed with liquid medium containing CN inhibitors for 3, 6, 12 or 24 hours (red arrowheads), before testing for HS induced seizures. Flies were also tested 12 and 24 hours after drug withdrawal. **B-D.** Behavioral analysis of HS induced seizures. **B.** Flies were fed with 0.3 mM or 1 mM of CsA for 12 hours. Feeding with 1 mM CsA reduces seizures by ∼75% (p<0.0001). The effect of CsA treatment on HS-induced seizures shows a significant dose-dependent reduction in seizure occurrence (N=3 groups of >15 flies/treatment). **C.** Summary of all time points for CsA treatment (N=3 groups of 15-20 flies/treatment. 6 hours feeding: p=0.005; 12/24 hours feeding: p<0.0001; 12 hours drug withdrawal: p=0.0022). **D.** After 2 minutes of heat-shock, seizures were reduced by ∼50% (p=0.041) in flies that were fed with 0.3 mM CsA, and by ∼80% (p=0.0062) in flies that were fed with 1 mM CsA (N=3 groups of >15 flies/treatment). Error bars are SEM, *=P<0.05, **=P<0.01, ****=P<0.0001, Student’s t-test.

### Cortex glial knockdown of the sandman two-pore-domain K^+^ channel mimics zyd seizures

To explore how CN hyperactivation promotes seizures, we conducted a screen of known and putative CN targets using RNAi knockdown with repo-gal4. We concentrated our screen on putative CN target genes that are involved in signal transduction (Table S3). This screen revealed that pan-glial knockdown of sandman (sand), the *Drosophila* homolog of TRESK (KCNK18) and a member of the two-pore-domain K^+^ channel family (K_2P_), caused adult flies to undergo TS-induced seizures similar to *zyd* mutants (Fig. 5A). Vortex-induced seizures in repo>sand^RNAi^ were less severe than those observed in *zyd*, with only ∼50% of sand^RNAi^ flies showing seizures (Fig. S3A). TS-induced seizures in repo>sand^RNAi^ adults were found to have the same kinetics and temperature threshold as seizures observed in *zyd* mutants (Fig. 5A, B), and CPG recordings showed that repo>sand^RNAi^ larvae exhibit rapid, unpatterned firing at 38°C, similar to *zyd* mutants (Fig. 5C). Cortex-glial specific knockdown of sand recapitulated ∼50% of the seizure effect when two copies of the RNAi were expressed (Fig. 5A, B). The less robust effect observed with the cortex-glial driver could be due to less effective RNAi knockdown or secondary to a role for sand in other glial subtypes. To determine if sand functions in other glia subtypes to mimic the *zyd* seizure pathway, we expressed sand^RNAi^ using the pan-glial driver repo-gal4 and inhibited expression specifically in cortex glia with GMR54H02>gal80. In the absence of cortex glial-knockdown of sand, seizure generation was suppressed (Fig. 5A). Similar to *zyd* mutants, sand^RNAi^ animals did not show changes in general activity and locomotion at room temperature (Fig. 5D). Finally, RNAi knockdown of sand did not enhance the *zyd* phenotype (Fig. S3B), suggesting seizures due to loss of sand and zyd impinge on a similar pathway.

**Figure 5.**
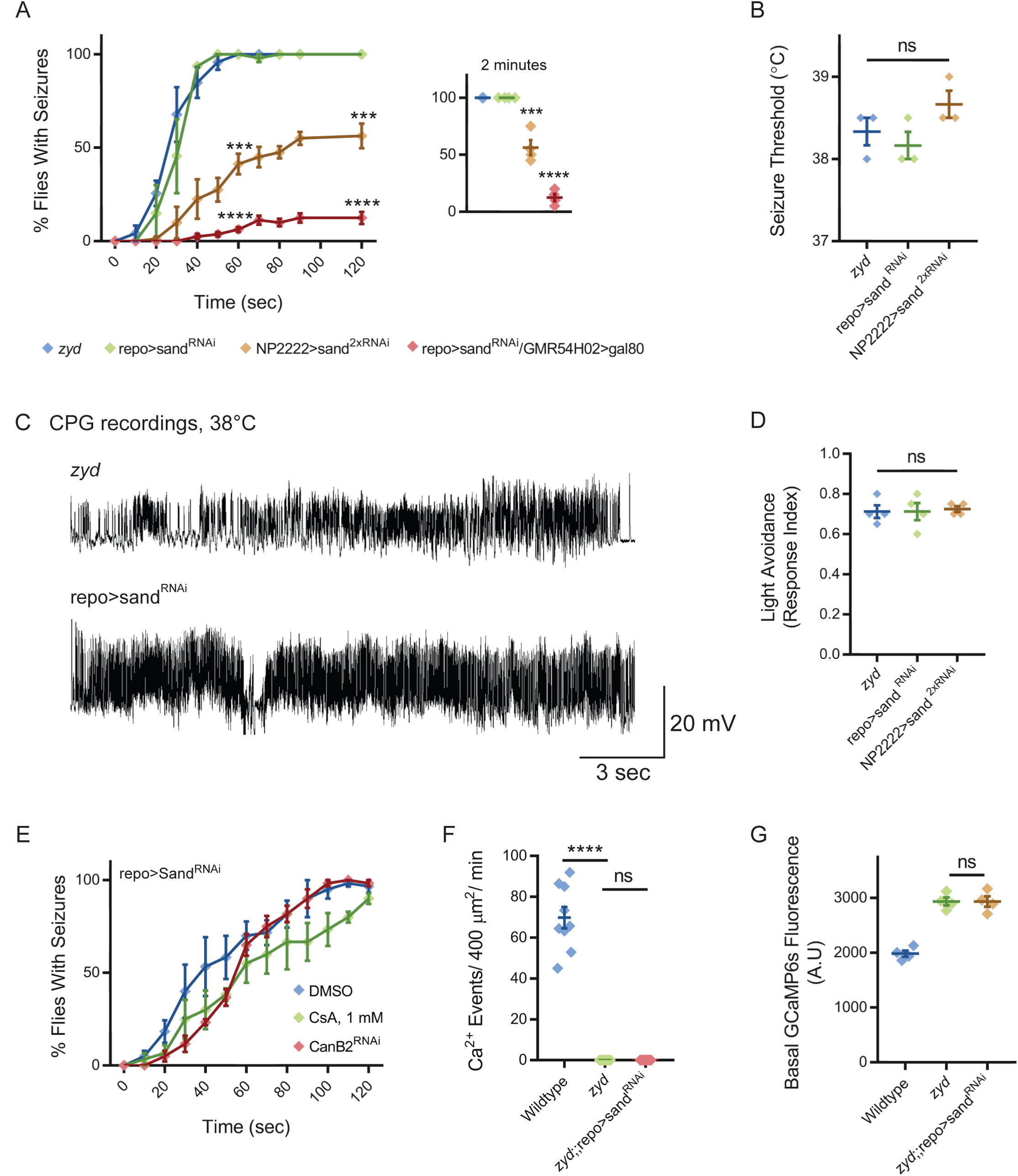
Cortex glial knock-down of sandman, a K_2P_ channel, reproduces *zyd* phenotypes. **A-B.** Behavioral analysis of HS induced seizures. **A.** Knockdown of sandman (sand) in different glial subtypes: pan-glial (repo), cortex glial (NP2222) and in all glia other than cortex glia (repo>sand^RNAi^/GMR54H02>gal80 in which gal80 is constitutively inhibiting gal4 activity and sand^RNAi^ expression occurs only in cortex glia). Inset shows analysis after 2 minutes of HS (p=0.0006 for NP2222>sand^2xRNAi^, p<0.0001 for repo>sand^RNAi^/GMR54H02-gal80, N=4 groups of >10 flies/genotype). **B.** Temperature threshold of repo-sand^RNAi^ (p=0.5185) and NP2222>sand^2xRNAi^ (p=0.2302) seizures in comparison to *zyd* (N=3 groups of 10/temperature/genotype). **C.** Representative voltage traces of spontaneous CPG activity at larval 3^rd^ instar muscle 6 at 38°C in *zyd* and repo>sand^RNAi^ (*n*≥5 preparations/genotype). **D.** Light avoidance assay reveals no defect in this behavior at 25°C (N=3 groups of 20 flies/genotype). **E.** Behavioral analysis of HS-induced seizures. Seizures in repo>sand^RNAi^ animals were not suppressed with either CanB2^RNAi#1^ or by feeding flies with 1 mM (N=3 groups of 20 flies/treatment). **F-G.** Ca^2+^ imaging in larval cortex glial cells using myrGCaMP6s. **F.** The average rate of microdomain Ca^2+^ events was reduced in repo>sand^RNAi^ cortex glia relative to wildtype (20.36 ± 5.5 and 69.83 ± 5.3, p<0.0001). Knockdown of sand on the *zyd* background did not restore *zyd* Ca^2+^ microdomain events (n≥5 animals/genotype). **G.** Average myrGCaMP6s fluorescence in cortex glia at 25°C. Elevated basal fluorescence of GCaMP6s in *zyd* relative to wildtype cortex glia (p=0.0003) is not altered following sand knockdown (*zyd*;;repo>sand^RNAi^, N=4 animals/genotype). Error bars are SEM, *=P<0.05, **=P<0.01, ***=P<0.001, ****=P<0.0001, Student’s t-test.

Mammalian astrocytes modulate neuronal network activity through regulation of K^+^ buffering (Bellot-Saez et al., 2017), in addition to their role in uptake of neurotransmitters such as GABA and glutamate (Murphy-Royal et al., 2017). Human K_ir4.1_ potassium channels (*KCNJ10*) have been implicated in maintaining K^+^ homeostasis, with mutations in the loci causing epilepsy (Haj-Yasein et al., 2011). In addition, gain and loss of astrocytic K_ir4.1_ influence the burst firing rate of neurons through astrocyte-to-neuronal cell body contacts (Cui et al., 2018). However, K_ir_ channels are unlikely to be the only mechanism for glial K^+^ clearance, as K_ir4.1_ channels account for less than half of the K^+^ buffering capacity of mature hippocampal astrocytes (Ma et al., 2014). To determine if cortex glial K_ir_ channels regulate seizure susceptibility in addition to sand, we used repo-gal4 to knock down all three *Drosophila* K_ir_ family members (Irk1, Irk2 and Irk3). Pan-glial knockdown of the *Drosophila* K_ir_ family did not cause seizures (Fig. S3C), while knock down of either Irk1 or Irk2 slightly enhanced the *zyd* phenotype (Fig. S3D). Similarly, repo-gal4 knockdown of other well-known *Drosophila* K^+^ channels beyond the K_ir_ family also did not cause seizures (Fig. S4C), indicating sand is likely to play a preferential role in K^+^ buffering in *Drosophila* cortex glia.

The mammalian sand homolog, TRESK, is directly activated by CN dephosphorylation (Czirjak et al., 2004; Enyedi and Czirjak, 2015), while *Drosophila* sand was shown to be modulated in sleep neurons by activity-induced internalization from the plasma membrane (Pimentel et al., 2016). Regardless of the mechanism by which CN may regulate the protein, we hypothesized that sand is epistatic to CN in controlling *zyd*-mediated seizures. Indeed, inhibition of CN by RNAi or CsA did not alter sand^RNAi^-induced seizures (Fig. 5E), placing sand downstream of CN activity. Furthermore, knockdown of sand in the *zyd* background does not alter the elevated basal Ca^2+^ or the lack of microdomain Ca^2+^ events in *zyd* mutants (Fig. 5F, G), suggesting sand is downstream to the abnormal Ca^2+^ signaling in *zyd.* Overall, these findings suggest elevated Ca^2+^ in *zyd* mutants leads to hyperactivation of CN and subsequent reduction in sand function. These results suggest that impairment in glial buffering of the rising extracellular K^+^ during elevated neuronal activity and stress conditions (i.e. heat shock or acute vortex) causes enhanced seizure susceptibility in *zyd* mutants.

### Enhanced endocytosis in zyd cortex glia

We next sought to examine how elevated CN activity in *zyd* mutants alters sand function. The mammalian sand homolog, TRESK, is constitutively phosphorylated on four serine residues (S264 by PKA, and S274, S276 and S279 by MARK1) (Enyedi and Czirjak, 2015). Two of these residues are conserved in *Drosophila* sand (S264 and S276, see Fig. S3E for protein alignment). Constitutive dephosphorylation and subsequent activation of sand by CN would be predicted to increase K^+^ buffering following hyperactivation of the nervous system by stressors, and thus less seizure activity would be expected – opposite to what we have observed. If this regulatory mechanism is active in *Drosophila* cortex glia, we hypothesized that knockdown of either PKA or Par-1 (the *Drosophila* MARK1 homolog) would lead to enhanced activity of sand and improvement or rescue of *zyd* seizures. However, neither pan-glial or cortex glial knockdown of either kinase (PKA-C1, PKA-C2, PKA-C3 and Par-1), or overexpression of a PKA inhibitory peptide (PKI), altered the *zyd* phenotype (Table S4). Together with the prediction that regulation of sand by dephosphorylation should lead to seizure suppression, these results argue against enhanced sand dephosphorylation as the primary cause of *zyd* seizures.

A second mechanism to link CN hyperactivation to sand regulation is suggested by a previous study demonstrating sand expression on the plasma membrane of neurons involved in sleep homeostasis is regulated by activity-dependent internalization (Pimentel et al., 2016). Cam and CN activate several endocytic Ca^2+^ sensors and effectors that control Ca^2+^-dependent endocytosis (Xie et al., 2017). If hyperactivity of CN leads to enhanced internalization of sand and subsequent seizure susceptibility due to decreased K^+^ buffering capacity, interrupting cortex glial endocytosis should suppress *zyd* seizures. To test this model, we used cortex glial-specific RNAi to knock down genes involved in endocytosis and early endosomal processing and trafficking. Cortex glial knockdown of several essential endocytosis genes, including dynamin-1 and clathrin heavy and light chains, caused embryonic lethality (Fig. S2A). In contrast, cortex glial knockdowns of Rab5, a Rab GTPase regulator of early endosome (EE) dynamics (Dunst et al., 2015; Langemeyer et al., 2018), and Endophilin A (EndoA), a BAR-domain protein involved in early stages of endocytosis (Verstreken et al., 2002), were found to be viable (Fig. S2A) and to completely suppress *zyd* TS seizures (Fig. 6A). A second, non-overlapping hairpin and a dominant negative (DN) construct for Rab5 (Rab5^DN^) resulted in early larval lethality (Table S5), likely due to more efficient Rab5 activity suppression. Cortex glial knockdown of Rab5 was previously shown to cause a morphological defect in which cortex glia fail to infiltrate the cortex and enwrap neuronal cell bodies (Coutinho-Budd et al., 2017). The loss of neuronal wrapping by cortex glia in Rab5^RNAi^ might influence the glia-to-neuron signaling pathway that is activated in *zyd* animals to increase their seizure susceptibility. To test whether the rescue effect of Rab5 knockdown is due to an impairment in the structure of glial-neuronal contacts, we conditionally expressed Rab5^RNAi^ and Rab5^DN^ both in 3^rd^ instar larvae and in adult cortex glia. Adult flies incubated for 16 hours at the restrictive temperature for gal80^ts^ (31°c) to allow Rab5^RNAi^ expression, showed a partial rescue in *zyd* seizures, with only ∼25% of flies showing HS induced seizures (Fig. 6B). Adult flies and 3^rd^ instar larvae conditionally expressing Rab5^DN^, showed a partial rescue in *zyd* seizures with only ∼40% of animals showing wildtype-like behavior (Fig. 6B). These results suggest that Rab5, a master regulator of endocytosis and EE biogenesis plays a key role for the pathway that is activated in *zyd* cortex glia to promote seizures (see below). However, it is also possible that the observed Rab5^RNAi^ rescue resulted from a general disruption in cellular membrane trafficking. To test this possibility, we assayed seizure suppression in *zyd* mutants following knockdown of the entire family of *Drosophila* Rab GTPases (Table S5), most of which are expressed in *Drosophila* cortex glia (Coutinho-Budd et al., 2017). Beyond Rab5, none of the Rab knockdowns altered the *zyd* seizure phenotype or caused a behavioral phenotype on a wildtype background (Table S5). To assay if excess endocytosis secondary to CN hyperactivity disrupts membrane trafficking, we imaged endosomal compartments by over-expressing GFP-tagged Rab proteins in cortex glial cells (with GMR54H02-gal4) in control and *zyd* animals. We found that large (>0.1 μm^2^) Rab5-positive early endosomes accumulated in *zyd* cortex glia compared to controls (Fig. 6C). Feeding *zyd* larvae the CN inhibitor, CsA (1 mM), restored the number of Rab5 compartments to control levels (Fig. 6D). These results indicate CN hyperactivation secondary to elevated Ca^2+^ levels in *zyd* mutants increases endocytosis and the formation and accumulation of early endosomes in cortex glia.

**Figure 6.**
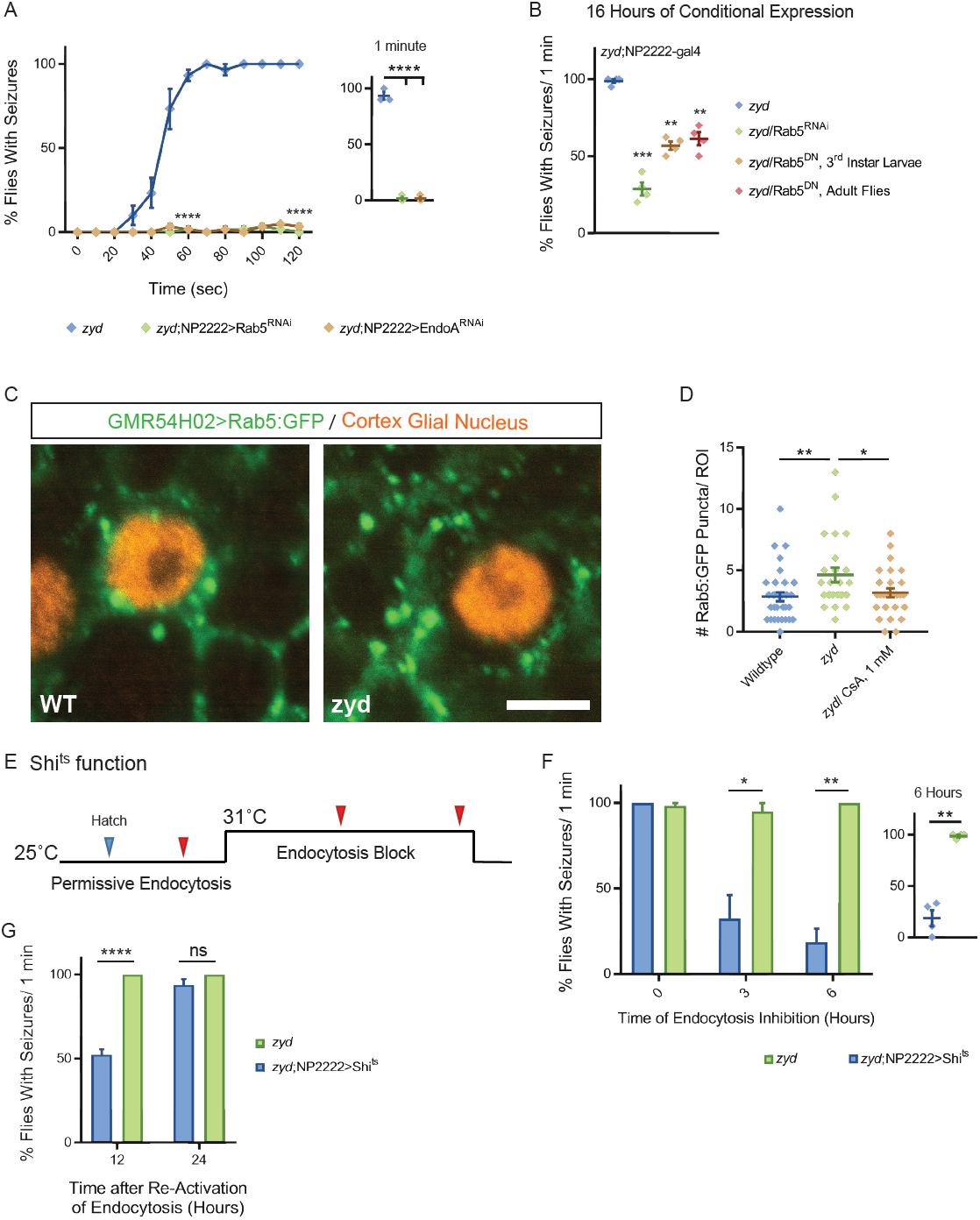
Cortex glial inhibition of endocytosis rescues *zyd* seizures. **A-B.** Behavioral analysis of HS induced seizures. **A.** Cortex glial knockdown of Rab5 and EndoA rescues seizures in the *zyd* mutant. Inset shows effects at 1 minute of HS (N=3 groups of 20 flies/genotype, p<0.0001 at 1 minute and 2 minutes). **B.** Cortex glial conditional overexpression of Rab5^RNAi^ and dominant-negative Rab5 (Rab5^DN^) using UAS/gal4/gal80^ts^. Rearing adult flies at the restrictive temperature (>30°C) for gal80^ts^ allows expression of Rab5^RNAi^ or Rab5^DN^ only in adults. A significant reduction in seizures was seen after 16 hours of incubation at the restrictive temperature for gal80^ts^ (31°C), with only ∼25% zyd/Rab5^RNAi^ flies showing seizures (p=0.0006), and ∼40% zyd/Rab5^DN^ larvae (p=0.0015)/ adult flies (p=0.0055) not showing seizures. (N=3 groups of >10 flies/genotype). **C.** Fluorescence images showing accumulation of Rab5::GFP puncta in *zyd* cortex glia relative to wildtype cortex glia. Rab5::GFP was expressed using a cortex glial-specific driver (GMR54H02-gal4. Scale bar=5 μm. n≥5 animals/genotype). **D.** Analysis of the number of large (>0.1 μm^2^) Rab5::GFP puncta in wildtype and *zyd* cortex glia. The number of Rab5::GFP puncta in *zyd* cortex glia was increased relative to wildtype (average of 4.64 ± 0.58 and 2.85 ± 0.37 puncta/ROI respectively, p=0.0088). The number of large Rab5::GFP puncta in *zyd* treated with 1 mM CsA for 24 hours was decreased relative to *zyd* (average of 3.19 ± 0.37 puncta/ROI, p=0.0378. n≥25 ROIs/3 animals/genotype/treatment). **E-F.** Conditional inhibition of endocytosis by cortex glial overexpression of shi^ts^. **E.** Schematic representation of the experimental design. Adult *zyd*; NP2222>shi^ts^ male flies (>1 day old) were incubated at the shi^ts^ restrictive temperature (31°C) for 3 or 6 hours and then tested for HS-induced seizures (red arrowheads, N=3 groups of >15 flies/time point). **F-G.** Behavioral analysis of HS induced seizures. **F.** A significant reduction in seizures is observed in flies that were incubated at 31°C for 3 hours (p=0.0283) or 6 hours (p=0.013). Inset shows analysis after 6 hours of incubation at 31°C (N=3 groups of >15 flies/time point). **G.** *zyd*;NP2222>Shi^ts^ flies seizures re-occur after removal from the Shi^ts^ restrictive temperature (N=4 groups of 10-15 animals/time point; 12 hours: p<0.0001). Error bars are SEM, *=P<0.05, **=P<0.01, ***=P<0.001, ****=P<0.0001, Student’s t-test.

### Chronic inhibition of dynamin-mediated endocytosis rescues zyd seizures

The partial rescue of the *zyd* seizure phenotype by conditional expression of Rab5^RNAi^ and Rab5^DN^ does not exclude the possibility that the rescue is partially due to morphological changes that may occur in these animals and not solely due to impairment in endocytosis. In addition, our previous analysis of the *zyd* mutant indicated that basal intracellular Ca^2+^ is elevated in cortex glia, with Ca^2+^ levels increasing even more when *zyd* animals are heat-shocked (Melom and Littleton, 2013). This additional elevation in Ca^2+^ could potentially further enhance CN activity and endocytosis beyond that observed at rest. These data raise two questions: 1) whether the basal enhancement of endocytosis or the additional heat shock-induced Ca^2+^ increase is the primary cause for seizure susceptibility in *zyd* mutants, and 2) Whether the rescue effect of Rab5^RNAi^ and EndoA^RNAi^ is due to changes in glia-to-neuron contacts morphology. To directly asses the role of endocytosis in increasing seizure susceptibility in *zyd* flies, we conditionally manipulated endocytosis by overexpressing a TS dominant-negative form of Dynamin-1 (Shi^ts^) in *zyd* cortex glia. This mutant version of Dynamin has normal activity at room temperature and a dominant-negative function upon exposure to the non-permissive temperature (>29°C, Fig. 6F). Acute inhibition of endocytosis by inactivation of Shi^ts^ in cortex glia did not suppress *zyd* seizures, suggesting further enhancement of CN activity and endocytosis specifically during the heat shock is not likely to be the cause of the rapid-onset seizures observed in *zyd* mutants. Given the chronic enhancement in CN activity and endocytosis in *zyd* mutants demonstrated by enhanced CalexA signaling (Fig. 3) and early endosome accumulation (Fig. 6C-D), we hypothesized that inhibiting endocytosis prior to exposing animals to a heat shock might improve their phenotype by altering the plasma membrane protein content over longer timescales. We incubated *zyd*;NP2222>Shi^ts^ flies at a non-permissive temperature for Shi^ts^ (31°C) for either 3 or 6 hours, and then tested for heat shock-induced seizures at 38.5°C (Fig. 6E-F). *Zyd* mutants alone do not undergo seizures at 31°C, nor does pre-incubation at 31°C alter the subsequent seizure phenotype observed at 38.5°C. In contrast, inhibition of endocytosis for 6 hours at 31°C in *zyd* mutants co-expressing Shi^ts^ suppressed the subsequent seizures observed during a 38.5°C heat shock in ∼80% of animals (Fig. 6F). A shorter 3 hours inhibition did not cause a significant improvement in seizures. The seizure suppression observed after 6 hours of Dynamin inhibition was reversible, as adults tested 12 or 24 hours after return to room temperature regained the *zyd* seizure phenotype (Fig. 6G). We conclude that chronic hyperactivation of CN and endocytosis caused by elevated basal Ca^2+^ in *zyd* cortex glia is the primary cause for *zyd* seizures.

### Increasing glial K^+^ uptake rescues zyd-dependent seizures

Genetic analysis of the *zyd* mutant indicate the primary cause of seizure susceptibility is chronic enhancement in Ca^2+^-dependent CN activity and subsequent increases in endocytosis in cortex glia. We hypothesize this enhancement in endocytosis leads to increased internalization of plasma membrane proteins such as sand, which in turn disrupt K^+^ uptake and buffering by cortex glial cells during periods of intense activation of the nervous system (Fig. 7A). To test this model further, we assayed if artificially increasing cortex glial K^+^ uptake in *zyd* mutants by overexpressing another K^+^ leak channel could suppress the seizure phenotype. Indeed, constitutive cortex glial overexpression of the open K^+^ channel EKO (White et al., 2001) rescued vortex-induced seizures in ∼75% of *zyd* mutants (Fig 7B). During a heat shock, cortex glial overexpression of EKO led to a dramatic change in the behavior of ∼60% of *zyd* animals, changing the seizure phenotype to hypoactivity or complete paralysis (Fig. 7C). CPG recordings revealed that *zyd*;;repo>EKO larvae lose all bursting and normal CPG activity at 38°C (Fig. 7D, middle), while *zyd*;NP2222>EKO regain normal wildtype-like rhythmic activity (Fig. 7D, right). These results indicate cortex glial K^+^ buffering is critical for neuronal excitability during states of intense excitation as observed following heat shock or acute vortex. During these periods of intense neuronal activity in *zyd* mutants, defective cortex glial K^+^ buffering surrounding neuronal somas leads to seizures. Enhancing K^+^ buffering can either reverse the seizure phenotype or push the scale toward neuronal hypo-excitability and paralysis. Future studies will be required to determine if sand function is dynamically modulated by normal microdomain Ca^2+^ oscillatory activity in wildtype cortex glia in response to changes in neuronal activity, which would provide a robust glial-based homeostatic mechanism to maintain neuronal spiking rates in acceptable ranges.

**Figure 7:**
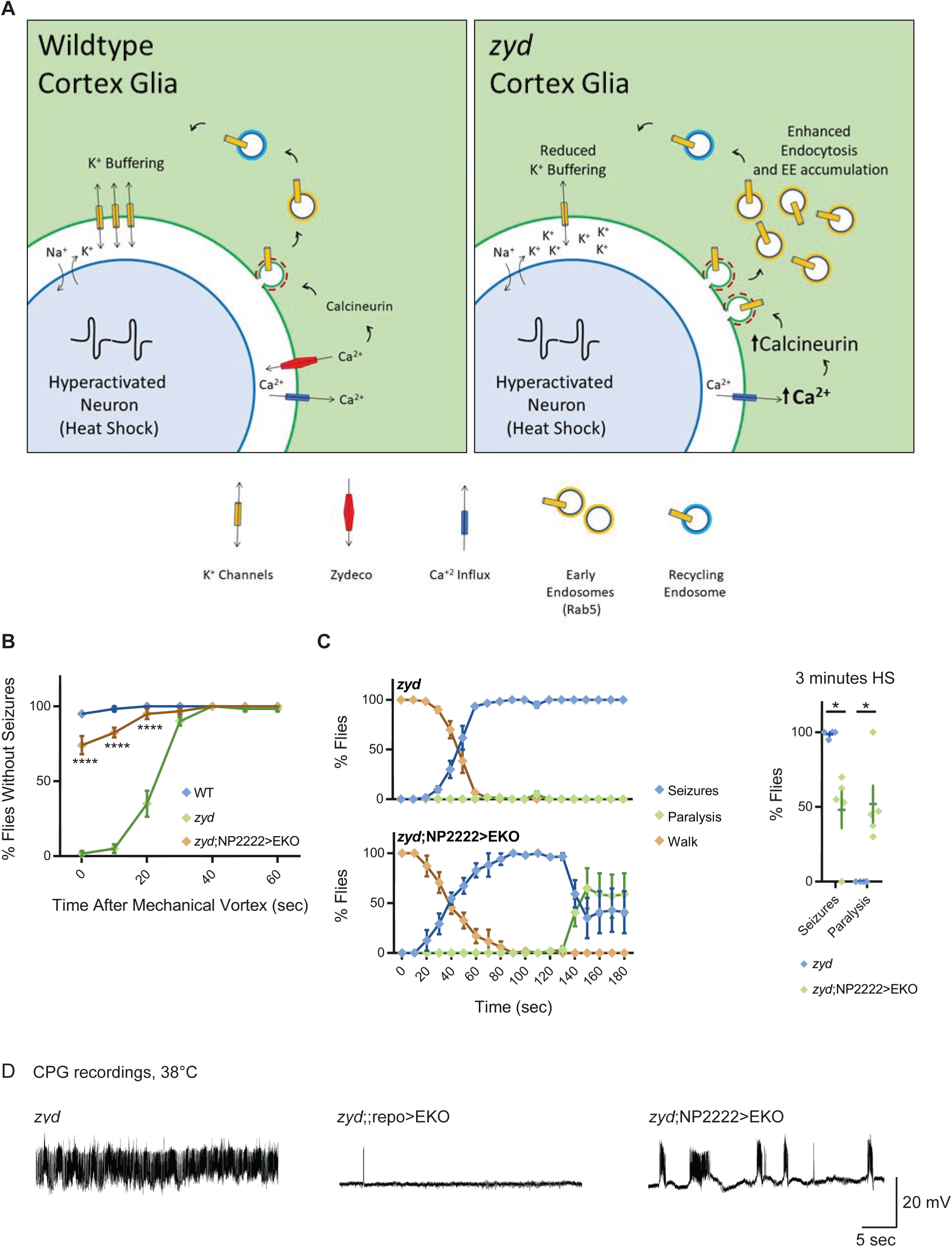
Enhancing glial K^+^ buffering by overexpressing a leak K^+^ channel rescues *zyd* seizures. **A.** The current model for *zyd* function in seizure susceptibility is depicted. In wildtype (WT) cortex glia (left), oscillatory Ca^2+^ signaling maintains normal cortex glia-to-neuron communication and a balanced extracellular ionic environment. In *zyd* cortex glia (right), the basal elevation of Ca^2+^ leads to hyperactivation of CN and enhanced endocytosis with accumulation of early endosomes. This disrupts the endo-exocytosis balance of cortex glial membrane proteins such as the K_2P_ leak channel sandman that regulate neuronal excitability. Knockdown of sandman mimics the *zyd* phenotype and the protein acts downstream of CN, indicating impaired extracellular K^+^ homeostasis causes neuronal hyperexcitability and hyperactivity in *zyd* mutants. **B-D.** Overexpression of a genetically modified constitutively-open Shaker K^+^ channel (termed EKO) modifies the *zyd* phenotype. **B.** *zyd* vortex-induced seizures are suppressed by ∼80% in *zyd*; NP2222>EKO flies relative to *zyd* alone (p<0.0001, N=3 groups of 20 flies/genotype). **C.** During a HS, *zyd*; NP2222>EKO flies transition from early seizures to hypoactivity and paralysis. Inset shows analysis after 3 minutes of HS (seizures: p=0.009; paralysis: p=0.008, N=5 groups of 20 flies/genotype). **D.** Representative voltage traces of spontaneous CPG activity at 3^rd^ instar larval muscle 6 at 38°C in *zyd*, *zyd*;;repo>EKO and *zyd*;NP2222>EKO (*n*≥3 preparations/genotype). EKO expression in all glia or only cortex glia eliminates the continuous CPG seizures observed in *zyd* mutants. Error bars are SEM, *=P<0.05, ****=P<0.0001, Student’s t-test.

### Discussion

Significant progress has been made in understanding glial-neuronal communication at synaptic and axonal contacts, but whether glia regulate neuronal function via signaling at somatic regions remains largely unstudied. A single mammalian astrocyte can be associated with multiple neuronal cell bodies and thousands of synapses, making it challenging to direct manipulations that alter glial signaling only at neuronal somas. *Drosophila* cortex glia provide an ideal system to explore how glia regulate neuronal function at the soma, as they ensheath multiple neuronal cell bodies, but do not contact synapses (Awasaki et al., 2008). In this study, we took advantage of the *zydeco* TS seizure mutation in a NCKX Ca^2+^ exchanger to explore pathways by which cortex glia regulate neuronal excitability. We found that elevation of basal Ca^2+^ levels in cortex glia leads to hyperactivation of Ca^2+^-CN dependent endocytosis. We showed that seizures in *zyd* mutants can be fully suppressed by either conditional inhibition of endocytosis or by pharmacologically reducing CN activity. Two well-characterized mechanisms by which glia regulate neuronal excitability and seizure susceptibility are neurotransmitter uptake via surface transporters and spatial K^+^ buffering. Cortex glia do not contact synapses, making it unlikely they are exposed to neurotransmitters. Instead, cortex glial-knockdown of the two-pore K^+^ channel (K_2P_) sandman, the *Drosophila* homolog of TRESK/KCNK18, recapitulates *zyd* TS seizures. These findings suggest impairment in K^+^ buffering during hyperactivity in *zyd* mutants underlies the increased seizure susceptibility, providing an unexpected link between glial Ca^2+^ signaling and K^+^ buffering. Consistent with this model (Fig. 7A), enhancing cortex glial K^+^ uptake by overexpressing a constitutively open K^+^ channel (EKO (White et al., 2001)) reduces neuronal activity in *zyd* mutants, rescues vortex-induced seizures, and reverses the TS behavioral phenotype from seizures to paralysis.

Different glial subtypes, including astrocytes, exhibit dynamic fluctuations in intracellular Ca^2+^ *in vitro* (Fatatis and Russell, 1992; Nett et al., 2002) and *in vivo* (Nimmerjahn et al., 2009; Porter and McCarthy, 1996). These early discoveries led to the idea that astrocytes can listen to and regulates neuronal and brain activity. Accumulated findings indicate that glial Ca^2+^ signaling influences neuronal physiology on a rapid time scale: increased glial Ca^2+^ activity is associated with abnormal neuronal excitability (Wetherington et al., 2008), and pathologic elevation of glial Ca^2+^ can play an important role in the generation of seizures (Gomez-Gonzalo et al., 2010; Tian et al., 2005). However, the signaling pathways underlying these astrocytic Ca^2+^ transients and their relevance to brain activity are poorly defined and controversial (Fiacco and McCarthy, 2018; Savtchouk and Volterra, 2018).

Previous work studying astrocytic Ca^2+^ signaling artificially increased intracellular Ca^2+^ using different approaches (i.e. pharmacological or optogenetic stimulation, caged Ca^2+^ photolysis and transgenic receptor expression) (Savtchouk and Volterra, 2018). However, whether these manipulations mimic physiological astrocytic responses is unclear. The *Drosophila zyd* mutation was identified in an unbiased genetic screen for behavioral mutants that triggered TS-dependent seizures, thus establishing the biological importance of the pathway before the gene mutation and cellular origin of the defect was known. The elevation in cortex glial Ca^2+^ levels found in *zyd* mutants provides a mechanism to explore how this pathway influences neuronal excitability. We recently found that Ca^2+^ elevation in astrocyte-like glia results in the rapid internalization of the astrocytic plasma membrane GABA transporter GAT and subsequent silencing of neuronal activity through elevation in synaptic GABA levels (Zhang et al., 2017). As such, Ca^2+^-regulated endo/exocytic trafficking of neurotransmitter transporters and K^+^ channels to and from the plasma membrane may represent a broadly used mechanism for linking glial Ca^2+^ activity to the control of neuronal excitability at synapses and cell bodies, respectively.

CN is the only Ca^2+^/Cam-dependent phosphatase encoded in the genome and is highly expressed throughout the brain (Furman and Norris, 2014; Goto et al., 1986a, b; Kuno et al., 1992; Polli et al., 1991). CN interacts with numerous neuronal substrates to modulate diverse functions, including receptor and ion channel trafficking, ion channel function and gene regulation (Baumgartel and Mansuy, 2012). Neuronal CN also controls synapse loss, dendritic atrophy, synaptic dysfunction, and neuronal vulnerability (Abdul et al., 2010; Reese and Taglialatela, 2011). In contrast, astrocytic CN expression has been shown to increase during aging, injury and disease. Glial cells rely on CN signaling pathways to regulate phenotypic switching/cellular activation, immune-like responses and cytokine production (Furman and Norris, 2014), with the pathway being intimately involved in neuroinflammation (Furman and Norris, 2014; Kataoka et al., 2009; Nagamoto-Combs and Combs, 2010; Rojanathammanee et al., 2013; Shiratori et al., 2010). The role of CN in astrocytes during normal brain states is unclear (Chen et al., 2016). The identification of calmodulin (Melom and Littleton, 2013) and CN as suppressors of glial-originated seizures in *zyd* mutants indicate a Cam/CN-dependent pathway is hyperactivated and impairs normal cortex glial-to-neuronal soma signaling. Although the mechanism by which CN activity is upregulated in *Drosophila* cortex glia is different from that observed in mammals, the ability of CN to alter neuronal activity appears similar to mechanisms employed in glial-dependent neuroinflammatory states. During injury or disease states in mammals, activated astrocytes exhibit Ca^2+^ dysregulation with higher intracellular Ca^2+^ levels, more frequent Ca^2+^ oscillations and an elevated expression of Ca^2+^ channels and Ca^2+^-regulated proteins (Kuchibhotla et al., 2009; Lin et al., 2007; Sama and Norris, 2013). While it is clear how these changes would hyperactivate CN signaling, which has extreme consequences for neuronal function, it is unclear what role basal CN activity has in glial signaling. CN regulates the expression of several key Ca^2+^ mediators in multiple cell types (Carafoli et al., 1999; Genazzani et al., 1999; Graef et al., 1999; Groth et al., 2007), including plasma membrane Ca^2+^ channels, intracellular Ca^2+^ release channels, and Ca^2+^-dependent enzymes (Baumgartel and Mansuy, 2012). However, no obvious phenotypes were observed when we decreased CN activity in CanB2^RNAi^ cortex glia in control animals (Fig. 3C), suggesting the primary function of CN is only engaged following states of Ca^2+^ elevation in cortex glia during periods of strong neuronal activity.

The link between elevated glial Ca^2+^ signaling and defects in K^+^ buffering was an unexpected observation in the *zyd* mutant. Effective removal of K^+^ from the extracellular space is vital for maintaining brain homeostasis and limits network hyperexcitability during normal brain function, as disruptions in K^+^ clearance have been linked to several pathological conditions (David et al., 2009; Leis et al., 2005; Somjen, 2002). In addition to ion homeostasis, astrocytic K^+^ buffering has been suggested as a mechanism for promoting hyperexcitability and engaging specific network activity (Bellot-Saez et al., 2017; Wang et al., 2012). Two mechanisms for astrocytic K^+^ clearance have been previously identified, including net K^+^ uptake (mediated by the Na^+^/K^+^ ATPase pump) and K^+^ spatial buffering (via passive K^+^ influx) (Bellot-Saez et al., 2017). While many studies indicate K_ir4.1_, a weakly inward rectifying K^+^ channel exclusively expressed in glial cells, as an important channel mediating astrocytic K^+^ buffering, it is unlikely to be the only mechanism for glial K^+^ clearance, as K_ir4.1_ channels account for less than half of the K^+^ buffering capacity of mature hippocampal astrocytes (Ma et al., 2014).

Human K_ir4.1_ potassium channels (KCNJ10) have been implicated in maintaining K^+^ homeostasis, with mutations in the loci causing several CNS pathologies including epilepsy and seizures (Bataveljic et al., 2012; Ferraro et al., 2004; Haj-Yasein et al., 2011; Kaiser et al., 2006; Magana et al., 2013; Scholl et al., 2009; Vit et al., 2008; Wilcock et al., 2009). In addition, several studies have linked members of the K_2P_ family, mainly TREK-1 and TWIK-1, to distinct aspects of astrocytic function (Olsen et al., 2015). TREK-1 was shown to regulate fast glutamate release by astrocytes (Woo et al., 2012), TWIK-1 and TREK-1 were shown to mediate passive K^+^ conductance in astrocytes (Hwang et al., 2014), and TWIK-1 was shown to be recruited to the astrocytic membrane by mGluR3 activation. Our finding that cortex glial knockdown of the *Drosophila* KCNK18/TRESK K_2P_ homolog sand triggers stress-induced seizures indicate glial sand is also involved in K^+^ homeostasis in the brain.

We considered two potential mechanisms for CN regulation of sand, including direct dephosphorylation and altered endocytic trafficking. At rest, the mammalian sand homolog TRESK is constitutively phosphorylated. Following generation of a Ca^2+^ signal, TRESK is dephosphorylated and activated by CN (Enyedi and Czirjak, 2015). Constitutive dephosphorylation and activation of sand by CN could potentially result in higher K^+^ efflux from cortex glia and neuronal depolarization, leading to higher seizure susceptibility. However, in this scenario, K^+^ buffering upon hyperactivation of the nervous system should be more efficient, and thus less seizures are expected. Cortex glia knockdown of the two kinases that phosphorylate TRESK, PKA and Par-1, did not cause seizures. As such, we found no evidence that sand activity is regulated by CN dephosphorylation of the protein directly in cortex glia.

Beyond dephosphorylation, sand expression on the plasma membrane of specific sleep homeostat neurons in *Drosophila* is regulated by activity-induced internalization (Pimentel et al., 2016). Given astrocytes can shape neuronal circuit activity by actively altering their K^+^ uptake capabilities (Wang et al., 2012), we propose that *Drosophila* cortex glia regulate the expression levels of sand (and potentially other K^+^ buffering proteins) on the cell membrane in a Ca^2+^-regulated fashion in normal animals. When Ca^2+^ is constitutively elevated in *zyd* mutants, this regulation is thrown out of balance. Indeed, inhibition of endocytosis in cortex glia can rescue *zyd* seizures, suggesting that a membrane protein responsible for the neuronal hyperexcitability phenotype is being abnormally internalized in *zyd* cortex glia. Bypassing sand function by improving cortex glial K^+^ buffering/uptake capacity through overexpression of a leak K^+^ channel (EKO (White et al., 2001)) can reverse the *zyd* phenotype from seizures (caused by neuronal hyperactivity) to paralysis (caused by neuronal hypoactivity). Together, these findings indicate elevated Ca^2+^ levels lead to hyperactivation of CN and elevated endocytosis, sand internalization, and impairment in K^+^ buffering by cortex glia in *zyd* mutant animals (Fig. 7A).

Accumulating evidence indicate glia play contributive or even causative roles in several neurological disorders, neurodevelopmental syndromes and neurodegenerative diseases including epilepsy, Fragile X syndrome and SMA respectively. Increased glial activity is associated with abnormal neuronal excitability (Wetherington et al., 2008), and pathologic elevation of glial Ca^2+^ has been suggested to play an important role in the generation of seizures (Gomez-Gonzalo et al., 2010; Tian et al., 2005). Approximately 50 million people worldwide have epilepsy, making it one of the most common neurological diseases globally (World Health Organization, 2018, http://www.who.int/en/). The traditional view assumes that epileptogenesis occurs exclusively in neurons. However, an astrocytic basis for epilepsy was proposed almost two decades ago (Gomez-Gonzalo et al., 2010; Tashiro et al., 2002; Tian et al., 2005). In a non-pathological state, glia display Ca^2+^ oscillations spontaneously (Takata and Hirase, 2008) and in response to physiological neuronal activity (Wang et al., 2006). One widely-studied output of Ca^2+^ oscillations is gliotransmission, the glial release of certain transmitters (Agulhon et al., 2008; Lee et al., 2010). Indeed, Ca^2+^-dependent glutamate release from astrocytes causes synchronous currents in neighboring neurons (Angulo et al., 2004; Fellin et al., 2006; Fellin et al., 2004), and is capable of eliciting action potentials (Pirttimaki et al., 2011), suggesting abnormally elevated glial Ca^2+^ may produce neuronal hypersynchrony through enhanced gliotransmission. In addition, *in vivo* work demonstrated that several anti-epileptic drugs reduce glial Ca^2+^ oscillations (Tian et al., 2005). Although increased glial activity has been associated with abnormal neuronal excitability, the role of glia in the development and maintenance of seizures, and the exact pathway(s) by which abnormal glial Ca^2+^ alter glia-to-neuron communication and neuronal excitability are poorly characterized (Wetherington et al., 2008). A second proposed mechanism by which astrocytes regulate neuronal excitability and seizure susceptibility involves the uptake and redistribution of K^+^ ions (Bellot-Saez et al., 2017; Wang et al., 2012) and neurotransmitters (Rose et al., 2017) from the extracellular space. In this study, we explored the pathways that are activated within glial cells in response to abnormally elevated glial Ca^2+^ that triggers seizures. We found that both mechanisms are at play in *Drosophila* cortex glial cells, with elevated intracellular Ca^2+^ leading to impaired K^+^ buffering. We identified several other suppressors of seizure induction in *zyd* mutants that are involved in GPCR signaling and vesicle trafficking, suggesting additional glial pathways may impact neuronal activity as well.

Current estimates suggest ∼70% of children and adults with epilepsy can be successfully treated with current anti-epileptic drugs (World Health Organization, 2018, http://www.who.int/en/). The observation that several anti-epileptic drugs reduce glial Ca^2+^ oscillations *in vivo* (Tian et al., 2005), together with the fact that ∼30% of epilepsy patients are non-responders, suggest that pharmacologically targeting glial pathways might be a promising avenue for future drug development in the field. Several neuronal seizure mutants in *Drosophila* have already been demonstrated to respond to common human anti-epileptic drugs, indicating key mechanisms that regulate neuronal excitability are conserved from *Drosophila* to humans. Indeed, *zyd*-induced seizures can be rescued when animals are fed a CN inhibitor (Fig. 4), indicating pharmacological targeting of the glial CN pathway can improve the outcome of a glial-derived seizure mutant. Prior studies have also shown improvement following treatment with the CN inhibitor FK506 in a rodent kindling model (Moia et al., 1994; Moriwaki et al., 1996). These data suggest CN activity regulates epileptogenesis in both *Drosophila* and mammalian models. Further characterization of how glia detect, respond, and actively shape neuronal excitability is critical to our understanding of neuronal communication and future development of new treatments for neurological diseases like epilepsy.

## Experimental Procedures

### Drosophila genetics and molecular biology

Flies were cultured on standard medium at 22°C unless otherwise noted. *zydeco* (*zyd*^1^*, here designated as zyd*) mutants were generated by ethane methyl sulfonate (EMS) mutagenesis and identified in a screen for temperature-sensitive (TS) behavioral phenotypes(Guan et al., 2005). The UAS/gal4 and LexAop/LexA systems were used to drive transgenes in glia, including repo-gal4(Lee and Jones, 2005), a pan-glial driver; NP2222-gal4, a cortex-glial specific driver; GMR54H02-gal4, expressed in a smaller set of cortex glial cells; and alrm-gal4, an astrocyte-like glial cell specific driver. The *UAS-dsRNAi* flies used in the study were obtained from the VDRC (Vienna, Austria) or the TRiP collection (Bloomington *Drosophila* Stock Center, Indiana University, Bloomington, IN, USA). All screened stocks are listed in supplementary material (Table S2). UAS-*myrGCaMP6s* was constructed by replacing GCaMP5 in the previously described myrGCaMP5 transgenic construct(Melom and Littleton, 2013). Transgenic flies were obtained by standard germline injection (BestGene Inc.). For all experiments described, only male larva and adults were used. In RNAi experiments, the animals also had the UAS-dicer2 transgenic element on the X chromosome to enhance RNAi efficiency. For survival assays, embryos were collected in groups of ∼50 and transferred to fresh vials (n=3). 3^rd^ instar larvae and/or pupae were counted. Survival rate (SR) was calculated as:

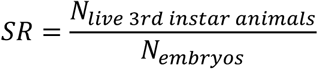

For conditional expression using Tub-gal80^ts^ (Figs. 1A, 2E, 6B and S3A), animals of the designated genotype were reared at 22°C with gal80 suppressing gal4-driven transgene expression (zyd^RNAi^, CanB2^RNAi^ and Rab5^DN^ and respectively). 3^rd^ instar larvae or adult flies were then transferred to a 31°C incubator to inactivate gal80 and allow gal4 expression or knockdown for the indicated period. For inhibiting transgene expression in cortex glia (Fig. 5A) GMR54H02-lexA (a kind gift from Gerald Rubin collection) was used to express gal80 from LexAop-gal80. For inhibiting transgene expression specifically in neurons (Fig. 2D) C155-gal80 was used.

Stocks used in this study:

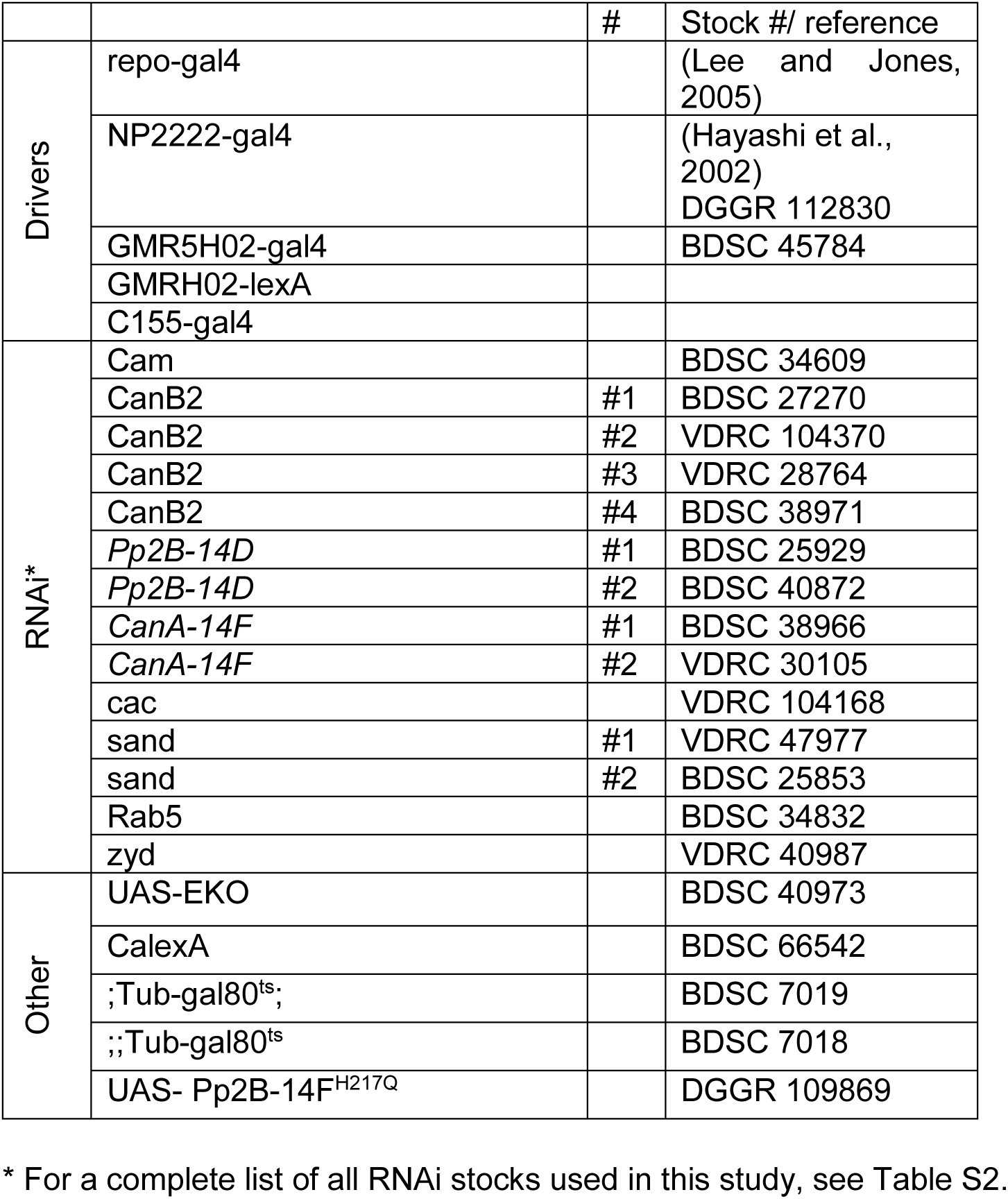

### Behavioral analysis

All experiments were performed using groups of ∼10-20 males.

#### 1. Temperature-sensitive seizures/zyd modifier screen

Adult males aged 1-2 days were transferred in groups of ∼10-20 flies (n≥3, total # of flies tested in all assays was always >40) into preheated vials in a water bath held at the indicated temperature with a precision of 0.1°C. Seizures were defined as the condition in which the animal lies incapacitated on its back or side with legs and wings contracting vigorously. Paralysis was defined as the condition in which the animal fell to the bottom of the vial and exhibited no movement. For screening purposes, only flies that showed normal wildtype-like behavior (i.e. walking up and down on vials walls) during heat-shock were counted as successful rescue. To analyze behavior in a more detailed manner we characterized four behavioral phenotypes: wall climbing (flies are climbing on vials walls), bottom dwelling (flies are on the bottom of the vial, standing/ walking without seizures), partial seizures (flies are on the bottom of the vial, seizing most of the time) and complete seizures (flies are constantly lying on their side or back with legs twitching). For assaying seizures in larvae, 3^rd^ instar males were gently washed with PBS and transferred to 1% agarose plates heated to 38°C using a temperature-controlled stage. Larval seizures were defined as continuous, unpatterned contraction of the body wall muscles that prevented normal crawling behavior. For determining seizure temperature threshold, groups of 10 animal were heat-shocked to the indicated temperature (either 37.5, 38, 38.5 or 39°C). Threshold was defined as the temperature in which >50% of the animal were seizing after 1 minute.

#### 2. Bang sensitivity

Adult male flies in groups of ∼10-20 males (n=3) were assayed 1-2 days post-eclosion. Flies were transferred into empty vials and allowed to rest for 1–2 h. Vials were vortexed at maximum speed for 10 seconds, and the number of flies that were upright and mobile was counted at 10 seconds intervals.

#### 3. Light avoidance

These assays were performed using protocols described previously following minor modifications. Briefly, pools of ∼20 3^rd^ instar larvae (108–120 hours after egg laying) were allowed to move freely for 5 minutes on Petri dishes with settings for the phototaxis assay (Petri dish lids were divided into quadrants, and two of these were blackened to create dark environment). The number of larvae in light versus dark quadrants was then scored (n=4). Response indices (RI) were calculated as:

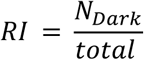

#### 4. Activity monitoring using the MB5 system

Adult flies activity was assayed using the multi-beam system (MB5, TriKinetics) as previously described (Green et al., 2015; McParland et al., 2015). Briefly, individual males aged 1-3 days were inserted into 5 mm×80 mm glass pyrex tubes. Activity was recorded following a 20-30 minutes acclimation period. Throughout each experiment, flies were housed in a temperature- and light-controlled incubator (25°C, ∼40-60% humidity). Post-acquisition activity analysis was performed using Excel to calculate activity level across 1-minute time bins (each experimental run contained 8 control animals and 8 experimental animals, n≥3).

#### 5. Gentle touch assay

3^rd^ instar male larvae (108–120 hours after egg laying) were touched on the thoracic segments with a hair during forward locomotion. No response, a stop, head retraction and turn were grouped into type I responses, and initiation of at least one single full body retraction or multiple full body retractions were categorized as type II reversal responses. Results were grouped to 20 males per assay (n=3).

### Electrophysiology

Intracellular recordings of wandering 3^rd^ instar male larvae were performed in HL3.1 saline (in mm: 70 NaCl, 5 KCl, 4 MgCl_2_, 0.2 CaCl_2_, 10 NaHCO_3_, 5 Trehalose, 115 sucrose, 5 HEPES-NaOH, pH 7.2) containing 1.5 mm Ca^2+^ using an Axoclamp 2B amplifier (Molecular Devices) at muscle fiber 6/7 of segments A3-A5. For recording the output of the central pattern generator, the CNS and motor neurons were left intact. Temperature was controlled with a Peltier heating device and continually monitored with a microprobe thermometer. Giant fiber recordings were performed as previously described (Pavlidis and Tanouye, 1995).

### *In vivo* Ca^2+^ imaging

UAS-*myrGCaMP6s* was expressed in glia *using the drivers described above.* 2^nd^ instar male larvae were washed with PBS and placed on a glass slide with a small amount of Halocarbon oil #700 (LabScientific). Larvae were turned ventral side up and gently pressed with a coverslip and a small iron ring to inhibit movement. Images were acquired with a PerkinElmer Ultraview Vox spinning disk confocal microscope and a high-speed EM CCD camera at 8–12 Hz with a 40X 1.3 NA oil-immersion objective using Volocity Software. Single optical planes within the ventral cortex of the ventral nerve cord (VNC) were imaged in the dense cortical glial region immediately below the surface glial sheath. Average myrGCaMP6s signal in cortex glia was quantified in the central abdominal neuromeres of the VNC within a manually selected ROI excluding the midline glia. Ca^2+^ oscillations were counted within the first minute of imaging at room temperature, and then normalized to the ROI area.

### Drug feeding

Cyclosporin-A (CsA, Sigma Aldrich) was dissolved in DMSO to a final concentration of 20 mM. Fly feeding solution included 5% yeast and 5% sucrose in water. Adult males less than 1 day old were starved for 6 hours and then transferred to a vial containing a strip of Wattman paper soaked in feeding solution containing the designated concentration of CsA or DMSO as control. Flies were behaviorally tested following 6, 12 or 24 hours of drug feeding.

### Immunostaining and Western Analysis

For immunostaining, dissected 3^rd^ instar male larvae were fixed with cold 4% paraformaldehyde in HL3.1 buffer for 45 minutes. Antibodies were used at the following dilutions: mouse anti-repo (8D12 Developmental Studies Hybridoma Bank), 1:50; rat anti-ELAV (7E8A10, Developmental Studies Hybridoma Bank), 1:50; GFP Rabbit IgG, Alexa Fluor^®^ 488 Conjugate (Thermo Fisher, 1:500); Goat anti mouse Alexa Fluor^®^ 405 Conjugate (Life technologies, 1:2000) and Goat anti-rat Alexa Fluor^®^ 555 Conjugate (Invitrogen, 1:2000). Larvae were mounted in VECTASHIELD (Vector Labs) and imaged on a Zeiss LSM800 confocal microscope with ZEN software (Carl Zeiss MicroImaging) with oil-immersion 63/1.4 NA objectives. Rab5::GFP was expressed specifically in cortex glia using GMR54H02-gal4 driver. GFP puncta (>0.1 μm^2^) were detected automatically within a set circular ROI (with r=5 μm, centroid in the center of the repo positive nucleus) using Volocity software. Western blotting of adult whole-head and larval brain lysates was performed using standard laboratory procedure. Nitrocellulose membranes were probed with rabbit anti-cleaved DCP1 (Cell Signaling, 1:1000) and rabbit anti-GFP (Abcam, 1:500). Equal loading was assayed using mouse anti-syx1A (1:1000). Primary antibodies were detected with Alexa Fluor 680-conjugated and 800-conjugated anti-rabbit and anti-mouse (Invitrogen, 1:3000). Western blots were visualized with an Odyssey infrared scanner (Li-Cor).

### Statistical analysis

No statistical methods were used to predetermine sample size. All *n* numbers represent biological replicates. Data were pooled from 2–3 independent experiments. Immunofluorescence experiments (Ca^2+^ imaging, CalexA expression and Rab5 puncta characterization) were randomized and blinded. P values are represented as *=P<0.05, **=P<0.01, ***=P<0.001, ****=P<0.0001. *P* < 0.05 was considered significant. All data are expressed as mean ± SEM.

## Acknowledgements

This work was supported by NIH grants NS40296 and MH104536 to J.T.L. We thank the Bloomington *Drosophila* Stock Center (NIH P40OD018537), the Vienna *Drosophila* RNAi Center, the Harvard TriP Project, the KYOTO Stock Center, Marc Freeman (Vollum Institute) and Gerald Rubin (Janelia Research Campus) for providing *Drosophila* strains, the Developmental Studies Hybridoma Bank for antisera, Mark Tanouye for help with GF recordings, and members of the Littleton lab for helpful discussions and comments on the manuscript.

## Author Contributions

S.W. performed the *zydeco* genetic interaction screen, confocal imaging, biochemistry, and behavioral analysis. J.E.M. assisted with the genetic modifier screen. K.G.O. and Y.V.Z. performed the electrophysiology. J.T.L supervised the project. S.W. and J.T.L. designed experiments and wrote the paper.

## Declaration of Interests

The authors declare no competing interests.

## Supplemental materials

### Supplemental figures

**Figure S1.**
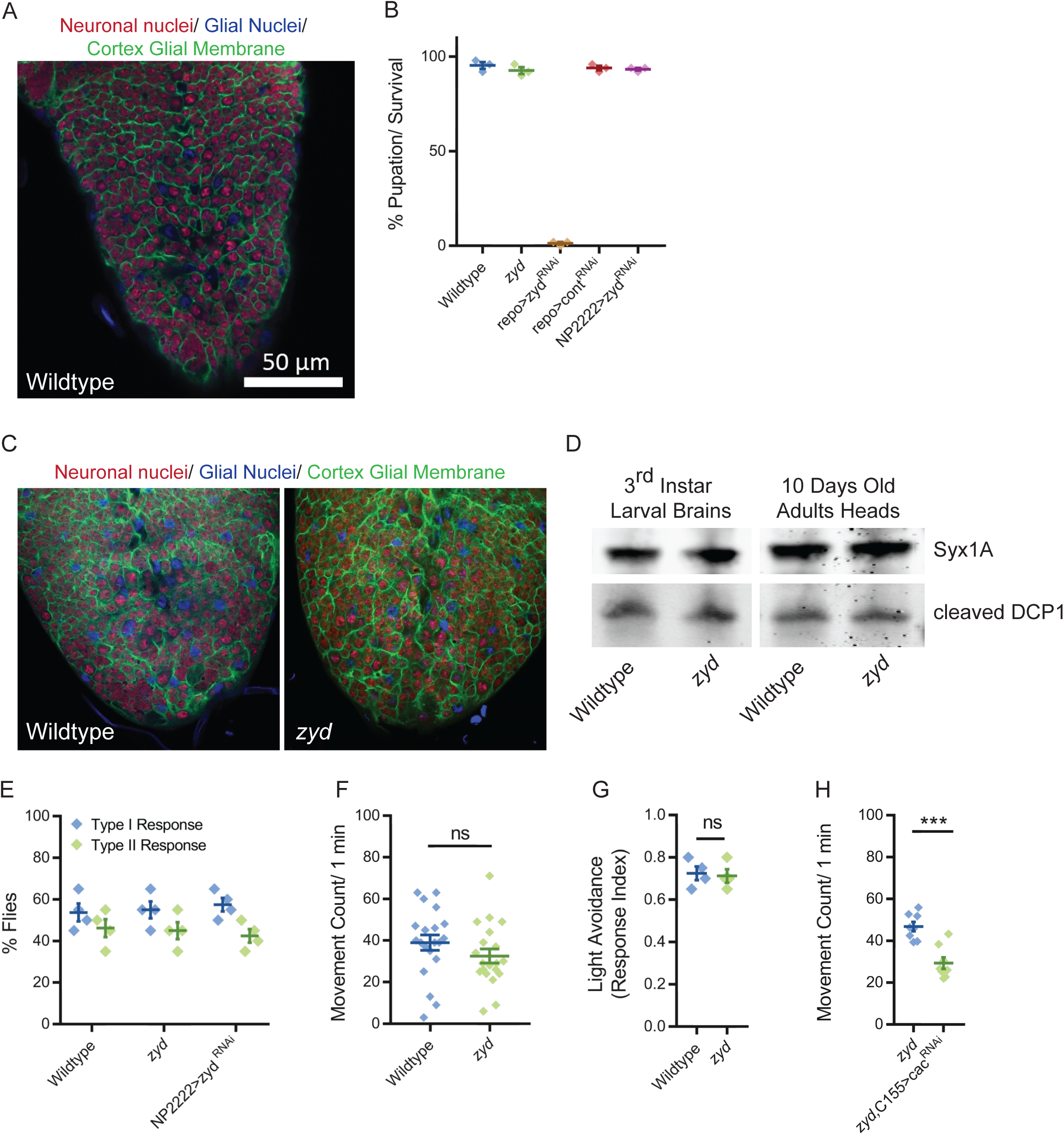
Mutations in a cortex glial NCKX generate stress-induced seizures without affecting brain structure or baseline neuronal function. *Related to Figure 1, Table S1 and Table S2.* **A.** Immunofluorescence imaging of a 3^rd^ instar larval ventral nerve cord (VNC) showing cortex glial membranes encapsulating neuronal soma (red: anti-Elav, neuronal nuclei; magenta: anti-repo, glial nuclei; green: anti-GFP, cortex glial membrane). **B.** Viability analysis of zyd^RNAi^ animals following expression of the RNAi with different glial drivers (N=3 groups of 50 embryos/genotype). Control was repo>GFP^RNAi^. **C.** Immunofluorescence imaging reveals no morphological changes in 3^rd^ instar larval brains associated with cortex glial wrapping of neuronal soma (red: anti-Elav, neuronal nuclei; magenta: anti-repo, glial nuclei; green: anti-GFP, cortex glial membrane). **D.** Western blot analysis of cleaved DCP1 (cell death marker) in larval brains and adult heads of wildtype and *zyd*. **E.** Behavioral analysis of the gentle touch response in wildtype and *zyd* larvae (N=4 groups of 20 larvae/genotype). **F.** Analysis of the activity level of wildtype and *zyd* adult flies (N=20 flies/genotype). **G.** Light avoidance response of wildtype and *zyd* 3^rd^ instar larvae (N=4 groups of 20 larvae/genotype). **H.** Activity level analysis shows a significant reduction in the activity of *zyd*/C155>cac^RNAi^ adult flies relative to *zyd* (p=0.0002) (n=8 flies/ genotype).

**Figure S2.**
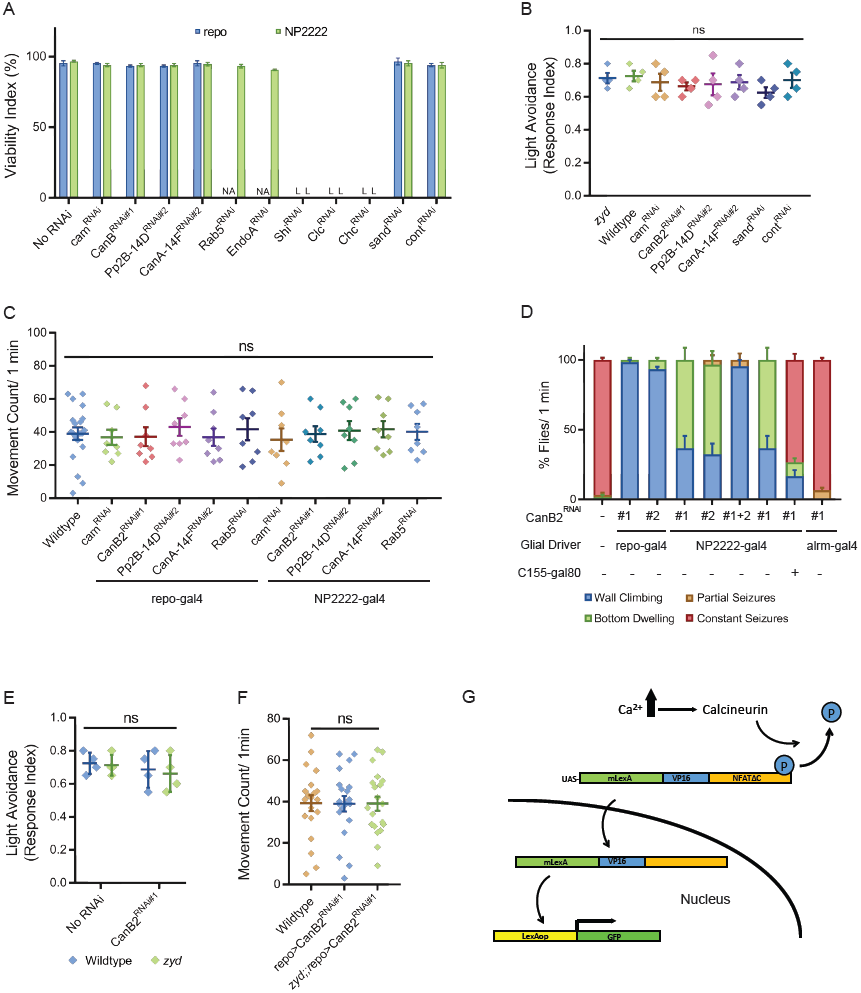
Cortex glial knockdown of calcineurin rescues *zyd* seizures without affecting intracellular Ca^2+^. *Related to Figure 2*-6. **A.** Viability analysis of the different RNAi used in this study. RNAi was expressed using a pan-glia (repo-gal4) or a cortex glial specific (NP-2222) driver (N=3 groups of 50 embryos/genotype). Control was GFP^RNAi^. **B.** Light avoidance response of 3^rd^ instar larvae expressing the different RNAi hairpins used in this study (N=4 groups of 20 larvae/genotype). **C.** Analysis of the activity level of flies expressing the different RNAi hairpins used in this study (N=8 flies/genotype). Control was GFP^RNAi^. **D.** Analysis of HS-induced behaviors of *zyd*/CanB2^RNAi^ flies (for full details, see methods). Cortex-glial knockdown of CanB2 leads to complete seizure rescue in ∼30% of *zyd*/CanB2^RNAi^ flies, while the remaining ∼70% show an intermediate phenotype. CanB2 knockdown with two copies of the RNAi recapitulates the more robust pan-glial knockdown effect. Inhibiting gal4 expression of the RNAi in neurons with gal80 (C155-gal80) does not alter the rescue effect seen in a single copy cortex glial knockdown (N=3 groups of >15 flies/genotype). **E.** Light avoidance response of wildtype, *zyd*, repo>CanB2^RNAi#1^ and *zyd*;;repo>CanB2^RNAi#1^ 3^rd^ instar larvae (N=4 groups of 20 larvae/genotype) **F.** Activity level analysis of wildtype, *zyd* and *zyd*;;repo>CanB2^RNAi#1^ adult flies (p=0.9843, n=20 flies/genotype). **G.** Schematic representation of the CalexA system. Sustained neural activity induces CN activation and dephosphorylation of a chimeric transcription factor LexA-VP16-NFAT (termed CalexA) which is then transported into the nucleus. The imported dephosphorylated CalexA drives GFP reporter expression in Ca^2+^-active cells.

**Figure S3.**
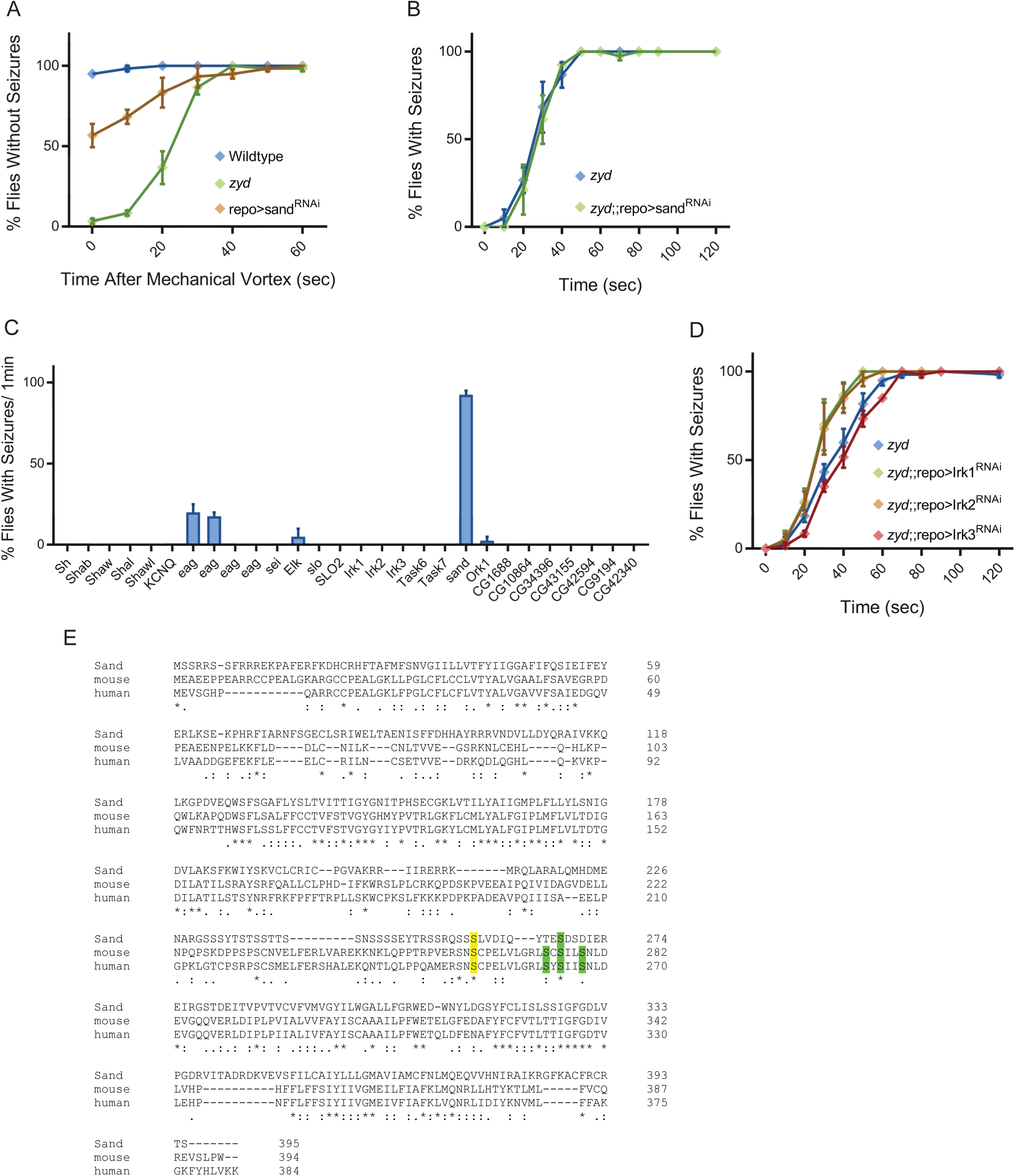
Cortex glial knockdown of sandman, a K_2P_ channel, reproduces *zyd* phenotypes. *Related to Figure 5 and Table S4.* **A.** Quantification of vortex-induced seizures in wildtype, *zyd* and repo>sand^RNAi^ adults. **B-C.** Behavioral analysis of HS-induced seizures. **B.** Pan-glial knockdown of sand on the *zyd* background does not enhance the *zyd* seizure phenotype (N=4 groups of >10 flies/genotype). **C.** Pan-glial knockdown of members of the *Drosophila* K_ir_ family does not enhance *zyd* seizures (N=4 groups of >10 flies/genotype). **C.** Pan-glial knockdown of other members of the *Drosophila* K^+^ channel family besides sand do not cause seizures (N=2 groups of ≥10 flies/genotype). **D.** Alignment of the protein sequences of mammalian KCNK18 and *Drosophila* SAND. Consensus phosphorylation sites for PKA (yellow) and MARK1 (green) are shown.

### Supplemental tables

**Table S1.**
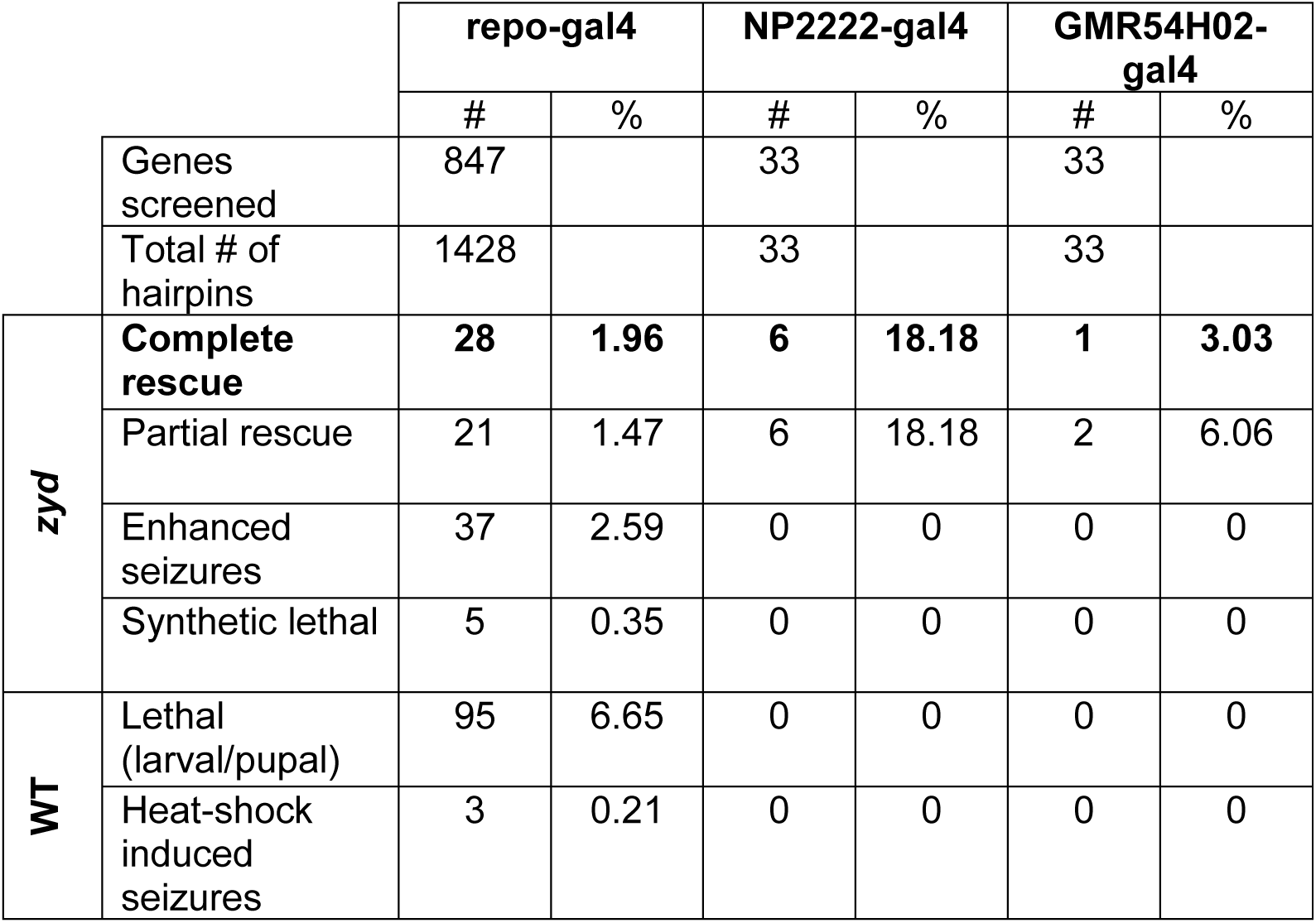
Summary of zyd suppressor/enhancer RNAi screen. *Related to Figure 1.*

|  |  | repo-gal4 |  | NP2222-gal4 |  | GMR54H02-gal4 |  |
| --- | --- | --- | --- | --- | --- | --- | --- |
|  |  | # | % | # | % | # | % |
| Genes screened |  | 847 |  | 33 |  | 33 |  |
| Total # of hairpins |  | 1428 |  | 33 |  | 33 |  |
| ***zyd*** | **Complete rescue** | **28** | **1.96** | **6** | **18.18** | **1** | **3.03** |
|  | Partial rescue | 21 | 1.47 | 6 | 18.18 | 2 | 6.06 |
|  | Enhanced seizures | 37 | 2.59 | 0 | 0 | 0 | 0 |
|  | Synthetic lethal | 5 | 0.35 | 0 | 0 | 0 | 0 |
| **WT** | Lethal (larval/pupal) | 95 | 6.65 | 0 | 0 | 0 | 0 |
|  | Heat-shock induced seizures | 3 | 0.21 | 0 | 0 | 0 | 0 |

**Tables S2 and S3 are in a supplemental Excel file**

**Table S4.** PKA/CamKII/Par-1 Transgenic Lines Assayed. *Related to Figure 5.*

| Gene | Stock # | repo>RNAi | zyd;;repo>RNAi | NP2222>RNAi | zyd;;NP2222>RNAi |
| --- | --- | --- | --- | --- | --- |
| **CaMKI** | 26726 | No phenotype | No phenotype | NA | NA |
|  | 35362 | No phenotype | No phenotype | NA | NA |
|  | 41900 | No phenotype | No phenotype | NA | NA |
| **CaMKII** | 29401 | No phenotype | Slightly enhanced seizures | No phenotype | No phenotype |
|  | v100265 | No phenotype | No phenotype | NA | NA |
|  | 35330 | No phenotype | No phenotype | NA | NA |
| **Par-1** | 32410 | No phenotype | No phenotype | No phenotype | No phenotype |
|  | 35342 | No phenotype | No phenotype | No phenotype | No phenotype |
| **Pka-C1** | BL31599 | No phenotype | No phenotype | NA | NA |
|  | BL31277 | No phenotype | No phenotype | NA | NA |
|  | BL35169 | No phenotype | No phenotype | NA | NA |
| **Pka-C2** | 31243 | No phenotype | No phenotype | NA | NA |
|  | 31656 | No phenotype | No phenotype | NA | NA |
|  | 55859 | No phenotype | No phenotype | NA | NA |
| **Pka-C3** | 27569 | No phenotype | No phenotype | NA | NA |
|  | 39050 | No phenotype | No phenotype | NA | NA |
|  | 55860 | No phenotype | No phenotype | NA | NA |
| **Pka-R1** | 27308 | No phenotype | No phenotype | NA | NA |
|  | 27708 | No phenotype | No phenotype | NA | NA |
|  | 52906 | No phenotype | No phenotype | NA | NA |
| **UAS-Pka-R1\* PKA inhibitor** | 35550 | No phenotype | No phenotype | NA | NA |

**Table S5.** Summary of Rab family knockdown in glia. *Related to Figure 6.* NS= no seizures; S= seizures; L= lethal

| gene | CG | RNAi | WT | *zyd* | WT | *zyd* | ctx glia expression<br>(Courtino-Budd, 2017) |
| --- | --- | --- | --- | --- | --- | --- | --- |
|  |  |  | repo-gal4 | repo-gal4 | NP222<br>2-gal4 | NP222<br>2-gal4 |  |
| Rab1 | CG3320 | JF02609 | L | L | L | L | Ubiquitous |
| Rab2 | CG3269 | JF03117 | NS | S | NS | S | Ubiquitous |
| Rab3 | CG7576 | JF02020 | NS | S | NS | S | Not detected |
| Rab3 | CG7576 | HMS01131 | NS | S | NS | S | Not detected |
| Rab4 | CG4921 | HMS01100 | NS | S | NS | S | Ubiquitous |
| Rab5 | CG3664 | JF03335 | L | L | L | L | Ubiquitous |
| **Rab5** | **CG3664** | **HMS00147** | **NS** | **NS** | **NS** | **NS** | **Ubiquitous** |
| Rab5 | CG3664 | Dominant<br>negative | L | L | L | L | Ubiquitous |
| Rab6 | CG6601 | JF02640 | NS | S | NS | S | Ubiquitous |
| Rab7 | CG5915 | JF02377 | NS | S | NS | S | Ubiquitous |
| Rab8 | CG8287 | JF02669 | NS | S | NS | S | Ubiquitous |
| Rab9 | CG9994 | JF01861 | NS | S | NS | S | Ubiquitous |
| Rab10 | CG17060 | JF02058 | NS | S | NS | S | Ubiquitous |
| Rab11 | CG5771 | JF02812 | NS | S | NS | S | Ubiquitous |
| Rab14 | CG4212 | JF03135 | NS | S | L | L | Not detected |
| Rab14 | CG4212 | HMS01130 | NS | S | NS | S | Not detected |
| Rab18 | CG3129 | JF02744 | NS | S | NS | S | Ubiquitous |
| Rab19 | CG7062 | HMS00592 | NS | S | NS | S | Ubiquitous |
| Rab21 | CG40304 | JF03338 | NS | S | NS | S | Ubiquitous |
| Rab23 | CG2108 | JF02859 | NS | S | NS | S | Ubiquitous |
| Rab26 | CG34410 | JF01684 | NS | S | NS | S | Ubiquitous |
| Rab27 | CG14791 | JF02165 | NS | S | NS | S | Ubiquitous |
| Rab30 | CG9100 | JF01593 | NS | S | NS | S | Not detected |
| Rab35 | CG9575 | JF02978 | NS | S | NS | S | Ubiquitous |
| Rab39 | CG12156 | JF01973 | NS | S | NS | S | Ubiquitous |
| Rab40 | CG1900 | JF03258 | NS | S | NS | S | Ubiquitous |
| RabX1 | CG3870 | JF02868 | NS | S | NS | S | Enriched |
| RabX2 | CG2885 | HMS00351 | NS | S | NS | S | Ubiquitous |
| RabX3 | CG32670 | HMS01364 | - | - | NS | S | Ubiquitous |
| RabX4 | CG13638 | JF03121 | NS | S | NS | S | Not detected |
| RabX5 | CG7980 | JF02880 | NS | S | NS | S | Ubiquitous |
| RabX6 | CG12015 | JF02050 | NS | S | NS | S | Ubiquitous |

## References

Abdul, H.M., Furman, J.L., Sama, M.A., Mathis, D.M., and Norris, C.M. (2010). NFATs and Alzheimer’s Disease. Mol Cell Pharmacol 2, 7–14.

Agulhon, C., Petravicz, J., McMullen, A.B., Sweger, E.J., Minton, S.K., Taves, S.R., Casper, K.B., Fiacco, T.A., and McCarthy, K.D. (2008). What is the role of astrocyte calcium in neurophysiology? Neuron 59, 932–946.

Akagawa, H., Hara, Y., Togane, Y., Iwabuchi, K., Hiraoka, T., and Tsujimura, H. (2015). The role of the effector caspases drICE and dcp-1 for cell death and corpse clearance in the developing optic lobe in Drosophila. Dev Biol 404, 61–75.

Allen, N.J., and Barres, B.A. (2009). Neuroscience: Glia - more than just brain glue. Nature 457, 675–677.

Altenhein, B., Becker, A., Busold, C., Beckmann, B., Hoheisel, J.D., and Technau, G.M. (2006). Expression profiling of glial genes during Drosophila embryogenesis. Dev Biol 296, 545–560.

Angulo, M.C., Kozlov, A.S., Charpak, S., and Audinat, E. (2004). Glutamate released from glial cells synchronizes neuronal activity in the hippocampus. J Neurosci 24, 6920–6927.

Awasaki, T., Lai, S.L., Ito, K., and Lee, T. (2008). Organization and postembryonic development of glial cells in the adult central brain of Drosophila. J Neurosci 28, 13742–13753.

Azevedo, F.A., Carvalho, L.R., Grinberg, L.T., Farfel, J.M., Ferretti, R.E., Leite, R.E., Jacob Filho, W., Lent, R., and Herculano-Houzel, S. (2009). Equal numbers of neuronal and nonneuronal cells make the human brain an isometrically scaled-up primate brain. J Comp Neurol 513, 532–541.

Baalman, K., Marin, M.A., Ho, T.S., Godoy, M., Cherian, L., Robertson, C., and Rasband, M.N. (2015). Axon initial segment-associated microglia. J Neurosci 35, 2283–2292.

Bataveljic, D., Nikolic, L., Milosevic, M., Todorovic, N., and Andjus, P.R. (2012). Changes in the astrocytic aquaporin-4 and inwardly rectifying potassium channel expression in the brain of the amyotrophic lateral sclerosis SOD1(G93A) rat model. Glia 60, 1991–2003.

Battefeld, A., Klooster, J., and Kole, M.H. (2016). Myelinating satellite oligodendrocytes are integrated in a glial syncytium constraining neuronal high-frequency activity. Nat Commun 7, 11298.

Baumgartel, K., and Mansuy, I.M. (2012). Neural functions of calcineurin in synaptic plasticity and memory. Learn Mem 19, 375–384.

Bellot-Saez, A., Kekesi, O., Morley, J.W., and Buskila, Y. (2017). Astrocytic modulation of neuronal excitability through K(+) spatial buffering. Neurosci Biobehav Rev 77, 87–97.

Buchanan, R.L., and Benzer, S. (1993). Defective glia in the Drosophila brain degeneration mutant drop-dead. Neuron 10, 839–850.

Carafoli, E., Genazzani, A., and Guerini, D. (1999). Calcium controls the transcription of its own transporters and channels in developing neurons. Biochem Biophys Res Commun 266, 624–632.

Chen, Y., Holstein, D.M., Aime, S., Bollo, M., and Lechleiter, J.D. (2016). Calcineurin beta protects brain after injury by activating the unfolded protein response. Neurobiol Dis 94, 139–156.

Cornell-Bell, A.H., Finkbeiner, S.M., Cooper, M.S., and Smith, S.J. (1990). Glutamate induces calcium waves in cultured astrocytes: long-range glial signaling. Science 247, 470–473.

Coutinho-Budd, J.C., Sheehan, A.E., and Freeman, M.R. (2017). The secreted neurotrophin Spatzle 3 promotes glial morphogenesis and supports neuronal survival and function. Genes Dev 31, 2023–2038.

Cui, Y., Yang, Y., Ni, Z., Dong, Y., Cai, G., Foncelle, A., Ma, S., Sang, K., Tang, S., Li, Y., et al. (2018). Astroglial Kir4.1 in the lateral habenula drives neuronal bursts in depression. Nature 554, 323–327.

Czirjak, G., Toth, Z.E., and Enyedi, P. (2004). The two-pore domain K+ channel, TRESK, is activated by the cytoplasmic calcium signal through calcineurin. J Biol Chem 279, 18550–18558.

Dani, J.W., Chernjavsky, A., and Smith, S.J. (1992). Neuronal activity triggers calcium waves in hippocampal astrocyte networks. Neuron 8, 429–440.

David, Y., Cacheaux, L.P., Ivens, S., Lapilover, E., Heinemann, U., Kaufer, D., and Friedman, A. (2009). Astrocytic dysfunction in epileptogenesis: consequence of altered potassium and glutamate homeostasis? J Neurosci 29, 10588–10599.

Ding, F., O’Donnell, J., Thrane, A.S., Zeppenfeld, D., Kang, H., Xie, L., Wang, F., and Nedergaard, M. (2013). alpha1-Adrenergic receptors mediate coordinated Ca2+ signaling of cortical astrocytes in awake, behaving mice. Cell Calcium 54, 387–394.

Dumstrei, K., Wang, F., and Hartenstein, V. (2003). Role of DE-cadherin in neuroblast proliferation, neural morphogenesis, and axon tract formation in Drosophila larval brain development. J Neurosci 23, 3325–3335.

Dunst, S., Kazimiers, T., von Zadow, F., Jambor, H., Sagner, A., Brankatschk, B., Mahmoud, A., Spannl, S., Tomancak, P., Eaton, S., et al. (2015). Endogenously tagged rab proteins: a resource to study membrane trafficking in Drosophila. Dev Cell 33, 351–365.

Enyedi, P., and Czirjak, G. (2015). Properties, regulation, pharmacology, and functions of the K(2)p channel, TRESK. Pflugers Arch 467, 945–958.

Fatatis, A., and Russell, J.T. (1992). Spontaneous changes in intracellular calcium concentration in type I astrocytes from rat cerebral cortex in primary culture. Glia 5, 95–104.

Fellin, T., Gomez-Gonzalo, M., Gobbo, S., Carmignoto, G., and Haydon, P.G. (2006). Astrocytic glutamate is not necessary for the generation of epileptiform neuronal activity in hippocampal slices. J Neurosci 26, 9312–9322.

Fellin, T., Pascual, O., Gobbo, S., Pozzan, T., Haydon, P.G., and Carmignoto, G. (2004). Neuronal synchrony mediated by astrocytic glutamate through activation of extrasynaptic NMDA receptors. Neuron 43, 729–743.

Ferraro, T.N., Golden, G.T., Smith, G.G., Martin, J.F., Lohoff, F.W., Gieringer, T.A., Zamboni, D., Schwebel, C.L., Press, D.M., Kratzer, S.O., et al. (2004). Fine mapping of a seizure susceptibility locus on mouse Chromosome 1: nomination of Kcnj10 as a causative gene. Mamm Genome 15, 239–251.

Fiacco, T.A., and McCarthy, K.D. (2018). Multiple Lines of Evidence Indicate That Gliotransmission Does Not Occur under Physiological Conditions. J Neurosci 38, 3–13.

Furman, J.L., and Norris, C.M. (2014). Calcineurin and glial signaling: neuroinflammation and beyond. J Neuroinflammation 11, 158.

Genazzani, A.A., Carafoli, E., and Guerini, D. (1999). Calcineurin controls inositol 1,4,5-trisphosphate type 1 receptor expression in neurons. Proc Natl Acad Sci U S A 96, 5797–5801.

Gomez-Gonzalo, M., Losi, G., Chiavegato, A., Zonta, M., Cammarota, M., Brondi, M., Vetri, F., Uva, L., Pozzan, T., de Curtis, M., et al. (2010). An excitatory loop with astrocytes contributes to drive neurons to seizure threshold. PLoS Biol 8, e1000352.

Goto, S., Matsukado, Y., Mihara, Y., Inoue, N., and Miyamoto, E. (1986a). Calcineurin in human brain and its relation to extrapyramidal system. Immunohistochemical study on postmortem human brains. Acta Neuropathol 72, 150–156.

Goto, S., Matsukado, Y., Mihara, Y., Inoue, N., and Miyamoto, E. (1986b). The distribution of calcineurin in rat brain by light and electron microscopic immunohistochemistry and enzyme-immunoassay. Brain Res 397, 161–172.

Graef, I.A., Mermelstein, P.G., Stankunas, K., Neilson, J.R., Deisseroth, K., Tsien, R.W., and Crabtree, G.R. (1999). L-type calcium channels and GSK-3 regulate the activity of NF-ATc4 in hippocampal neurons. Nature 401, 703–708.

Green, E.W., O’Callaghan, E.K., Pegoraro, M., Armstrong, J.D., Costa, R., and Kyriacou, C.P. (2015). Genetic analysis of Drosophila circadian behavior in seminatural conditions. Methods Enzymol 551, 121–133.

Groth, R.D., Coicou, L.G., Mermelstein, P.G., and Seybold, V.S. (2007). Neurotrophin activation of NFAT-dependent transcription contributes to the regulation of pro-nociceptive genes. J Neurochem 102, 1162–1174.

Guan, Z., Saraswati, S., Adolfsen, B., and Littleton, J.T. (2005). Genome-wide transcriptional changes associated with enhanced activity in the Drosophila nervous system. Neuron 48, 91–107.

Haj-Yasein, N.N., Jensen, V., Vindedal, G.F., Gundersen, G.A., Klungland, A., Ottersen, O.P., Hvalby, O., and Nagelhus, E.A. (2011). Evidence that compromised K+ spatial buffering contributes to the epileptogenic effect of mutations in the human Kir4.1 gene (KCNJ10). Glia 59, 1635–1642.

Halassa, M.M., Fellin, T., Takano, H., Dong, J.H., and Haydon, P.G. (2007). Synaptic islands defined by the territory of a single astrocyte. J Neurosci 27, 6473–6477.

Hayashi, S., Ito, K., Sado, Y., Taniguchi, M., Akimoto, A., Takeuchi, H., Aigaki, T., Matsuzaki, F., Nakagoshi, H., Tanimura, T., et al. (2002). GETDB, a database compiling expression patterns and molecular locations of a collection of Gal4 enhancer traps. Genesis 34, 58–61.

Hu, G., Wang, K., Groenendyk, J., Barakat, K., Mizianty, M.J., Ruan, J., Michalak, M., and Kurgan, L. (2014). Human structural proteome-wide characterization of Cyclosporine A targets. Bioinformatics 30, 3561–3566.

Hwang, E.M., Kim, E., Yarishkin, O., Woo, D.H., Han, K.S., Park, N., Bae, Y., Woo, J., Kim, D., Park, M., et al. (2014). A disulphide-linked heterodimer of TWIK-1 and TREK-1 mediates passive conductance in astrocytes. Nat Commun 5, 3227.

Kaiser, M., Maletzki, I., Hulsmann, S., Holtmann, B., Schulz-Schaeffer, W., Kirchhoff, F., Bahr, M., and Neusch, C. (2006). Progressive loss of a glial potassium channel (KCNJ10) in the spinal cord of the SOD1 (G93A) transgenic mouse model of amyotrophic lateral sclerosis. J Neurochem 99, 900–912.

Kataoka, A., Tozaki-Saitoh, H., Koga, Y., Tsuda, M., and Inoue, K. (2009). Activation of P2X7 receptors induces CCL3 production in microglial cells through transcription factor NFAT. J Neurochem 108, 115–125.

Kawasaki, F., Zou, B., Xu, X., and Ordway, R.W. (2004). Active zone localization of presynaptic calcium channels encoded by the cacophony locus of Drosophila. J Neurosci 24, 282–285.

Kuchibhotla, K.V., Lattarulo, C.R., Hyman, B.T., and Bacskai, B.J. (2009). Synchronous hyperactivity and intercellular calcium waves in astrocytes in Alzheimer mice. Science 323, 1211–1215.

Kuebler, D., and Tanouye, M. (2002). Anticonvulsant valproate reduces seizure-susceptibility in mutant Drosophila. Brain Res 958, 36–42.

Kuno, T., Mukai, H., Ito, A., Chang, C.D., Kishima, K., Saito, N., and Tanaka, C. (1992). Distinct cellular expression of calcineurin A alpha and A beta in rat brain. J Neurochem 58, 1643–1651.

Langemeyer, L., Frohlich, F., and Ungermann, C. (2018). Rab GTPase Function in Endosome and Lysosome Biogenesis. Trends Cell Biol.

Lee, B.P., and Jones, B.W. (2005). Transcriptional regulation of the Drosophila glial gene repo. Mech Dev 122, 849–862.

Lee, S., Bang, S.M., Hong, Y.K., Lee, J.H., Jeong, H., Park, S.H., Liu, Q.F., Lee, I.S., and Cho, K.S. (2016). The calcineurin inhibitor Sarah (Nebula) exacerbates Abeta42 phenotypes in a Drosophila model of Alzheimer’s disease. Dis Model Mech 9, 295–306.

Lee, S., Yoon, B.E., Berglund, K., Oh, S.J., Park, H., Shin, H.S., Augustine, G.J., and Lee, C.J. (2010). Channel-mediated tonic GABA release from glia. Science 330, 790–796.

Leis, J.A., Bekar, L.K., and Walz, W. (2005). Potassium homeostasis in the ischemic brain. Glia 50, 407–416.

Lin, D.T., Wu, J., Holstein, D., Upadhyay, G., Rourk, W., Muller, E., and Lechleiter, J.D. (2007). Ca2+ signaling, mitochondria and sensitivity to oxidative stress in aging astrocytes. Neurobiol Aging 28, 99–111.

Ma, B., Xu, G., Wang, W., Enyeart, J.J., and Zhou, M. (2014). Dual patch voltage clamp study of low membrane resistance astrocytes in situ. Mol Brain 7, 18.

Ma, Z., Stork, T., Bergles, D.E., and Freeman, M.R. (2016). Neuromodulators signal through astrocytes to alter neural circuit activity and behaviour. Nature 539, 428–432.

Magana, J.J., Velazquez-Perez, L., and Cisneros, B. (2013). Spinocerebellar ataxia type 2: clinical presentation, molecular mechanisms, and therapeutic perspectives. Mol Neurobiol 47, 90–104.

Masuyama, K., Zhang, Y., Rao, Y., and Wang, J.W. (2012). Mapping neural circuits with activity-dependent nuclear import of a transcription factor. J Neurogenet 26, 89–102.

McParland, A.L., Follansbee, T.L., and Ganter, G.K. (2015). Measurement of larval activity in the Drosophila activity monitor. J Vis Exp, e52684.

Melom, J.E., and Littleton, J.T. (2013). Mutation of a NCKX eliminates glial microdomain calcium oscillations and enhances seizure susceptibility. J Neurosci 33, 1169–1178.

Moia, L.J., Matsui, H., de Barros, G.A., Tomizawa, K., Miyamoto, K., Kuwata, Y., Tokuda, M., Itano, T., and Hatase, O. (1994). Immunosuppressants and calcineurin inhibitors, cyclosporin A and FK506, reversibly inhibit epileptogenesis in amygdaloid kindled rat. Brain Res 648, 337–341.

Moriwaki, A., Lu, Y.F., Hayashi, Y., Tomizawa, K., Tokuda, M., Itano, T., Hatase, O., and Matsui, H. (1996). Immunosuppressant FK506 prevents mossy fiber sprouting induced by kindling stimulation. Neurosci Res 25, 191–194.

Murphy-Royal, C., Dupuis, J., Groc, L., and Oliet, S.H.R. (2017). Astroglial glutamate transporters in the brain: Regulating neurotransmitter homeostasis and synaptic transmission. J Neurosci Res 95, 2140–2151.

Nagamoto-Combs, K., and Combs, C.K. (2010). Microglial phenotype is regulated by activity of the transcription factor, NFAT (nuclear factor of activated T cells). J Neurosci 30, 9641–9646.

Nakai, Y., Horiuchi, J., Tsuda, M., Takeo, S., Akahori, S., Matsuo, T., Kume, K., and Aigaki, T. (2011). Calcineurin and its regulator sra/DSCR1 are essential for sleep in Drosophila. J Neurosci 31, 12759–12766.

Nedergaard, M. (1994). Direct signaling from astrocytes to neurons in cultures of mammalian brain cells. Science 263, 1768–1771.

Nett, W.J., Oloff, S.H., and McCarthy, K.D. (2002). Hippocampal astrocytes in situ exhibit calcium oscillations that occur independent of neuronal activity. J Neurophysiol 87, 528–537.

Nimmerjahn, A., Mukamel, E.A., and Schnitzer, M.J. (2009). Motor behavior activates Bergmann glial networks. Neuron 62, 400–412.

Olsen, M.L., Khakh, B.S., Skatchkov, S.N., Zhou, M., Lee, C.J., and Rouach, N. (2015). New Insights on Astrocyte Ion Channels: Critical for Homeostasis and Neuron-Glia Signaling. J Neurosci 35, 13827–13835.

Parker, L., Howlett, I.C., Rusan, Z.M., and Tanouye, M.A. (2011). Seizure and epilepsy: studies of seizure disorders in Drosophila. Int Rev Neurobiol 99, 1–21.

Parpura, V., Basarsky, T.A., Liu, F., Jeftinija, K., Jeftinija, S., and Haydon, P.G. (1994). Glutamate-mediated astrocyte-neuron signalling. Nature 369, 744–747.

Paukert, M., Agarwal, A., Cha, J., Doze, V.A., Kang, J.U., and Bergles, D.E. (2014). Norepinephrine controls astroglial responsiveness to local circuit activity. Neuron 82, 1263–1270.

Pavlidis, P., and Tanouye, M.A. (1995). Seizures and failures in the giant fiber pathway of Drosophila bang-sensitive paralytic mutants. J Neurosci 15, 5810–5819.

Pimentel, D., Donlea, J.M., Talbot, C.B., Song, S.M., Thurston, A.J.F., and Miesenbock, G. (2016). Operation of a homeostatic sleep switch. Nature 536, 333–337.

Pirttimaki, T.M., Hall, S.D., and Parri, H.R. (2011). Sustained neuronal activity generated by glial plasticity. J Neurosci 31, 7637–7647.

Polli, J.W., Billingsley, M.L., and Kincaid, R.L. (1991). Expression of the calmodulin-dependent protein phosphatase, calcineurin, in rat brain: developmental patterns and the role of nigrostriatal innervation. Brain Res Dev Brain Res 63, 105–119.

Porter, J.T., and McCarthy, K.D. (1996). Hippocampal astrocytes in situ respond to glutamate released from synaptic terminals. J Neurosci 16, 5073–5081.

Reese, L.C., and Taglialatela, G. (2011). A role for calcineurin in Alzheimer’s disease. Curr Neuropharmacol 9, 685–692.

Rieckhof, G.E., Yoshihara, M., Guan, Z., and Littleton, J.T. (2003). Presynaptic N-type calcium channels regulate synaptic growth. J Biol Chem 278, 41099–41108.

Rojanathammanee, L., Puig, K.L., and Combs, C.K. (2013). Pomegranate polyphenols and extract inhibit nuclear factor of activated T-cell activity and microglial activation in vitro and in a transgenic mouse model of Alzheimer disease. J Nutr 143, 597–605.

Rose, C.R., Felix, L., Zeug, A., Dietrich, D., Reiner, A., and Henneberger, C. (2017). Astroglial Glutamate Signaling and Uptake in the Hippocampus. Front Mol Neurosci 10, 451.

Rusnak, F., and Mertz, P. (2000). Calcineurin: form and function. Physiol Rev 80, 1483–1521.

Sama, D.M., and Norris, C.M. (2013). Calcium dysregulation and neuroinflammation: discrete and integrated mechanisms for age-related synaptic dysfunction. Ageing Res Rev 12, 982–995.

Savtchouk, I., and Volterra, A. (2018). Gliotransmission: Beyond Black-and-White. J Neurosci 38, 14–25.

Scholl, U.I., Choi, M., Liu, T., Ramaekers, V.T., Hausler, M.G., Grimmer, J., Tobe, S.W., Farhi, A., Nelson-Williams, C., and Lifton, R.P. (2009). Seizures, sensorineural deafness, ataxia, mental retardation, and electrolyte imbalance (SeSAME syndrome) caused by mutations in KCNJ10. Proc Natl Acad Sci U S A 106, 5842–5847.

Shiratori, M., Tozaki-Saitoh, H., Yoshitake, M., Tsuda, M., and Inoue, K. (2010). P2X7 receptor activation induces CXCL2 production in microglia through NFAT and PKC/MAPK pathways. J Neurochem 114, 810–819.

Somjen, G.G. (2002). Ion regulation in the brain: implications for pathophysiology. Neuroscientist 8, 254–267.

Song, J., and Tanouye, M.A. (2008). From bench to drug: human seizure modeling using Drosophila. Prog Neurobiol 84, 182–191.

Spindler, S.R., Ortiz, I., Fung, S., Takashima, S., and Hartenstein, V. (2009). Drosophila cortex and neuropile glia influence secondary axon tract growth, pathfinding, and fasciculation in the developing larval brain. Dev Biol 334, 355–368.

Srinivasan, R., Huang, B.S., Venugopal, S., Johnston, A.D., Chai, H., Zeng, H., Golshani, P., and Khakh, B.S. (2015). Ca(2+) signaling in astrocytes from Ip3r2(-/-) mice in brain slices and during startle responses in vivo. Nat Neurosci 18, 708–717.

Takasaki, C., Yamasaki, M., Uchigashima, M., Konno, K., Yanagawa, Y., and Watanabe, M. (2010). Cytochemical and cytological properties of perineuronal oligodendrocytes in the mouse cortex. Eur J Neurosci 32, 1326–1336.

Takata, N., and Hirase, H. (2008). Cortical layer 1 and layer 2/3 astrocytes exhibit distinct calcium dynamics in vivo. PLoS One 3, e2525.

Takeo, S., Tsuda, M., Akahori, S., Matsuo, T., and Aigaki, T. (2006). The calcineurin regulator sra plays an essential role in female meiosis in Drosophila. Curr Biol 16, 1435–1440.

Tashiro, A., Goldberg, J., and Yuste, R. (2002). Calcium oscillations in neocortical astrocytes under epileptiform conditions. J Neurobiol 50, 45–55.

Tian, G.F., Azmi, H., Takano, T., Xu, Q., Peng, W., Lin, J., Oberheim, N., Lou, N., Wang, X., Zielke, H.R., et al. (2005). An astrocytic basis of epilepsy. Nat Med 11, 973–981.

Tomita, J., Mitsuyoshi, M., Ueno, T., Aso, Y., Tanimoto, H., Nakai, Y., Aigaki, T., Kume, S., and Kume, K. (2011). Pan-neuronal knockdown of calcineurin reduces sleep in the fruit fly, Drosophila melanogaster. J Neurosci 31, 13137–13146.

Ventura, R., and Harris, K.M. (1999). Three-dimensional relationships between hippocampal synapses and astrocytes. J Neurosci 19, 6897–6906.

Verstreken, P., Kjaerulff, O., Lloyd, T.E., Atkinson, R., Zhou, Y., Meinertzhagen, I.A., and Bellen, H.J. (2002). Endophilin mutations block clathrin-mediated endocytosis but not neurotransmitter release. Cell 109, 101–112.

Vit, J.P., Ohara, P.T., Bhargava, A., Kelley, K., and Jasmin, L. (2008). Silencing the Kir4.1 potassium channel subunit in satellite glial cells of the rat trigeminal ganglion results in pain-like behavior in the absence of nerve injury. J Neurosci 28, 4161–4171.

Volkenhoff, A., Weiler, A., Letzel, M., Stehling, M., Klambt, C., and Schirmeier, S. (2015). Glial Glycolysis Is Essential for Neuronal Survival in Drosophila. Cell Metab 22, 437–447.

Wang, F., Smith, N.A., Xu, Q., Fujita, T., Baba, A., Matsuda, T., Takano, T., Bekar, L., and Nedergaard, M. (2012). Astrocytes modulate neural network activity by Ca(2)+-dependent uptake of extracellular K+. Sci Signal 5, ra26.

Wang, X., Lou, N., Xu, Q., Tian, G.F., Peng, W.G., Han, X., Kang, J., Takano, T., and Nedergaard, M. (2006). Astrocytic Ca2+ signaling evoked by sensory stimulation in vivo. Nat Neurosci 9, 816–823.

Wetherington, J., Serrano, G., and Dingledine, R. (2008). Astrocytes in the epileptic brain. Neuron 58, 168–178.

White, B.H., Osterwalder, T.P., Yoon, K.S., Joiner, W.J., Whim, M.D., Kaczmarek, L.K., and Keshishian, H. (2001). Targeted attenuation of electrical activity in Drosophila using a genetically modified K(+) channel. Neuron 31, 699–711.

Wilcock, D.M., Vitek, M.P., and Colton, C.A. (2009). Vascular amyloid alters astrocytic water and potassium channels in mouse models and humans with Alzheimer’s disease. Neuroscience 159, 1055–1069.

Woo, D.H., Han, K.S., Shim, J.W., Yoon, B.E., Kim, E., Bae, J.Y., Oh, S.J., Hwang, E.M., Marmorstein, A.D., Bae, Y.C., et al. (2012). TREK-1 and Best1 channels mediate fast and slow glutamate release in astrocytes upon GPCR activation. Cell 151, 25–40.

Xie, Z., Long, J., Liu, J., Chai, Z., Kang, X., and Wang, C. (2017). Molecular Mechanisms for the Coupling of Endocytosis to Exocytosis in Neurons. Front Mol Neurosci 10, 47.

Zhang, Y.V., Ormerod, K.G., and Littleton, J.T. (2017). Astrocyte Ca(2+) Influx Negatively Regulates Neuronal Activity. eNeuro 4.

Zhou, Y., Cameron, S., Chang, W.T., and Rao, Y. (2012). Control of directional change after mechanical stimulation in Drosophila. Mol Brain 5, 39.

